# Do available protein 3D structures reflect human genetic and functional diversity?

**DOI:** 10.1101/637744

**Authors:** Gregory Sliwoski, Neel Patel, R. Michael Sivley, Charles R. Sanders, Jens Meiler, William S. Bush, John A. Capra

## Abstract

Genomic databases are substantially biased towards European ancestry populations, and this bias contributes to health disparities. Here, we quantify how well 66,971 experimentally characterized human protein 3D structures represent the diversity of protein sequences observed across the 1000 Genomes Project. More than 85% of available structures do not match a sequence observed in at least one individual, and on average structures match the sequence of 74% of individuals. Nearly 23% of human structures do not match *any* observed sequences; however, after masking engineered/known mutations, this decreases to ~4%. African ancestry sequences are modestly, but significantly, less likely to be represented by structures (73.5% vs. 74.0%). These differences are mainly driven by the greater genetic diversity of African populations. We identify thousands of variants unrepresented in available structures that influence protein structure and function. Thus, the use of a single structure as representative of “the wild type” protein will often bias results against many individuals. The diversity of protein sequence and structure must be considered to enable accurate, reproducible, and generalizable conclusions from structural analyses.

## INTRODUCTION

Genetic variants often influence human health and disease by modifying protein structure, stability, and function (de Beer et al., 2013; Gao et al., 2015; Wang and Moult, 2001). Protein structure is generally more strongly conserved than protein sequence (Chothia and Lesk, 1986); however, minor sequence changes can lead to dramatic structural changes (Barrera et al., 2016; Ben-Nissan et al., 2016; Bhattacharya et al., 2017; Creixell et al., 2015; Ferrer-Costa et al., 2002; Petukh et al., 2015). Advances in X-ray crystallography, nuclear magnetic resonance spectroscopy, electron microscopy (EM), and more recently cryo-EM have enabled modeling the 3D structure of thousands of human proteins and complexes (Berman et al., 2013; Ishchenko et al., 2018; Zeth and Zachariae, 2018). The Protein Data Bank (PDB, rcsb.org) (Berman et al., 2000)—the primary repository for experimentally determined protein structures—currently holds over 135,000 protein structures and is growing at a pace of approximately 10,000 structures per year. Analysis of 3D structures is a powerful and increasingly reliable tool in studies of the basic functions of proteins and how their dysfunction influences human disease (Blundell, 2017; Johansson and Lindorff-Larsen, 2018; Lu et al., 2013; Seeger, 2018; Sivley et al., 2018; Śledź and Caflisch, 2018). The broad importance of protein structures to basic and clinical research is demonstrated by the >160,000 citations of articles presenting human structures in PubMed Central (PMC), and the actual use of human structures is likely substantially higher due to the common practice of citing structures by their PDB ID without a primary reference.

Modern human populations exhibit substantial differences in levels of genetic diversity (1000 Genomes Project Consortium et al., 2015; Mallick et al., 2016; Mathias et al., 2016) and in the genetic architecture of many traits (Fan et al., 2016; Nielsen et al., 2017). As a result, studying the genetics of traits and diseases in diverse populations can yield greater power to map their genetic architectures and functional mechanisms (Crawford et al., 2017; Fumagalli et al., 2015). Despite the importance of considering diverse populations, our knowledge of genetic variation and its association with traits is strongly biased toward individuals of European ancestry (Popejoy and Fullerton, 2016). For example, more than 80% of genome-wide association studies (GWAS) have been performed in European ancestry populations, while African, Latin American and indigenous peoples of the Americas represent less than 4% of all individuals analyzed in GWAS (Carlson et al., 2013; Popejoy and Fullerton, 2016; Tekola-Ayele and Peprah, 2017). These and similar biases in clinical trials and genomic studies of rare diseases create substantial risk for genetic misdiagnoses and can perpetuate pre-existing health disparities (Hindorff et al., 2018). For example, African Americans are significantly more likely to receive a false positive diagnosis of hypertrophic cardiomyopathy based on genetic testing than white Americans due to failure to accurately capture patterns of genetic diversity among African Americans in genetic databases (Kessler et al., 2016; Manrai et al., 2016).

This problem is particularly challenging in the context of drug dosage, because clinically relevant pharmacogenomic loci commonly display allele frequency differences among human populations (Chan et al., 2016; Perera et al., 2014; Petrovski and Goldstein, 2016; Roden, 2016; Spooner et al., 2018; Tekola-Ayele et al., 2015). A recent examination of genetic variation in G-protein coupled receptors, one of the most important protein families for small molecule therapeutics, revealed that failure to account for genetic variation among individuals in drug prescription resulted in detrimental outcomes with an estimated financial impact of hundreds of millions of pounds a year in the UK alone (Hauser et al., 2018). In recognition of these challenges, the US NIH has prioritized inclusion of diverse populations in study of genetic variation and disease (Hindorff et al., 2018).

In light of the central role of protein structures in basic science, clinical analysis, and drug discovery, it is essential to evaluate the implications of human genetic diversity on protein structure and function. In this work, we quantify the extent to which available protein structures represent human genetic diversity and predict the effects of unrepresented common and rare genetic variants on protein structure. We find that available structures often represent common versions of a protein sequence, but rarely represent all individuals, and thus failure to account for human genetic diversity can result in inappropriate overgeneralization about protein function from structural analyses. Furthermore, we find that individuals of African ancestry are the least likely to be represented by and have the most sequence differences from available protein structures. However, these differences are less extreme than in many genomic databases. To help address these challenges, we quantify the degree to which each analyzed human protein structure represents different human populations and whether population-specific differences in sequence are likely to translate into altered structure and function.

## RESULTS

### Integrating population genetic variation and protein structures

In this study, we consider all single nucleotide missense variants identified in 2504 individuals from diverse human populations sequenced by the 1000 Genomes Project (1000G) across the 20,246 canonical protein isoforms in the Uniprot annotated human proteome (Apweiler et al., 2004). The 1000G individuals were sequenced using a combination of low-coverage whole genome sequencing (mean depth 7.4x) and deep exome sequencing (mean depth 65.7x). As expected from previous studies, most human protein isoforms exhibit sequence variation (1000 Genomes Project Consortium et al., 2010; Nelson et al., 2012; Tennessen et al., 2012) and do not have a single “wild type” sequence. The 1000G dataset contains 561,115 unique missense variants covering more than 87% (17,675 of 20,246) of human proteins (Table 1). Over 56% (11,414) of human proteins have at least one common missense variant (minor allele frequency (MAF) ≥ 0.01), and 365 proteins have >10 common variants (Figure S1). The average MAF is 0.13; however, the majority of missense variants (523,841) are rare (MAF < 0.01; Table 1, Figure S1).

**Table 1.** Number of missense variants in human proteins in the 1000G dataset stratified by minor allele frequency (MAF) and 1000G population. The number of proteins with variation (out of 20,246 total) is given in parentheses below the variant counts. AFR = African, AMR = Admixed American, EAS = East Asian, EUR = European, SAS = South Asian.

| VARIANTS | 1000G POPULATION |  |  |  |  |  |
| --- | --- | --- | --- | --- | --- | --- |
|  | ALL | AFR | AMR | EAS | EUR | SAS |
| **ALL** | 561,115<br>(17,675) | 217,635<br>(17,223) | 133,125<br>(16,565) | 141,599<br>(16,786) | 133,845<br>(16,727) | 150,874<br>(16,837) |
| **MAF $\geq 0.01$** | 37,274<br>(11,414) | 46,654<br>(12,440) | 31,457<br>(10,748) | 25,424<br>(9,785) | 28,589<br>(10,336) | 29,498<br>(10,424) |

To integrate this genetic variation data with protein structures, we mapped all missense variants into all available human protein structures from the PDB (31,681 unique PDB IDs) (Figure 1A). At the time of our analysis, the 66,971 human protein chains with structures in the PDB covered 27% (5,528 of 20,246) of human proteins (Figure 1B; Figure S1B). We will refer to each available quantification of the 3D location of atoms of an individual protein sequence as a “structure;” this corresponds to individual amino acid chains within PDB structures. Most proteins are represented by a small number of structures; however, 1,344 proteins have more than 10 structures available (Figure 1B). To account for multiple structures that overlap the same protein position, we performed all analyses over both the complete set of human protein structures (Main Text) and over a non-redundant set of structures selected to reflect protein coverage in the PDB while minimizing the amount of sequence position redundancy (Methods; Figure S2). Results were similar on each set.

**Figure 1.**
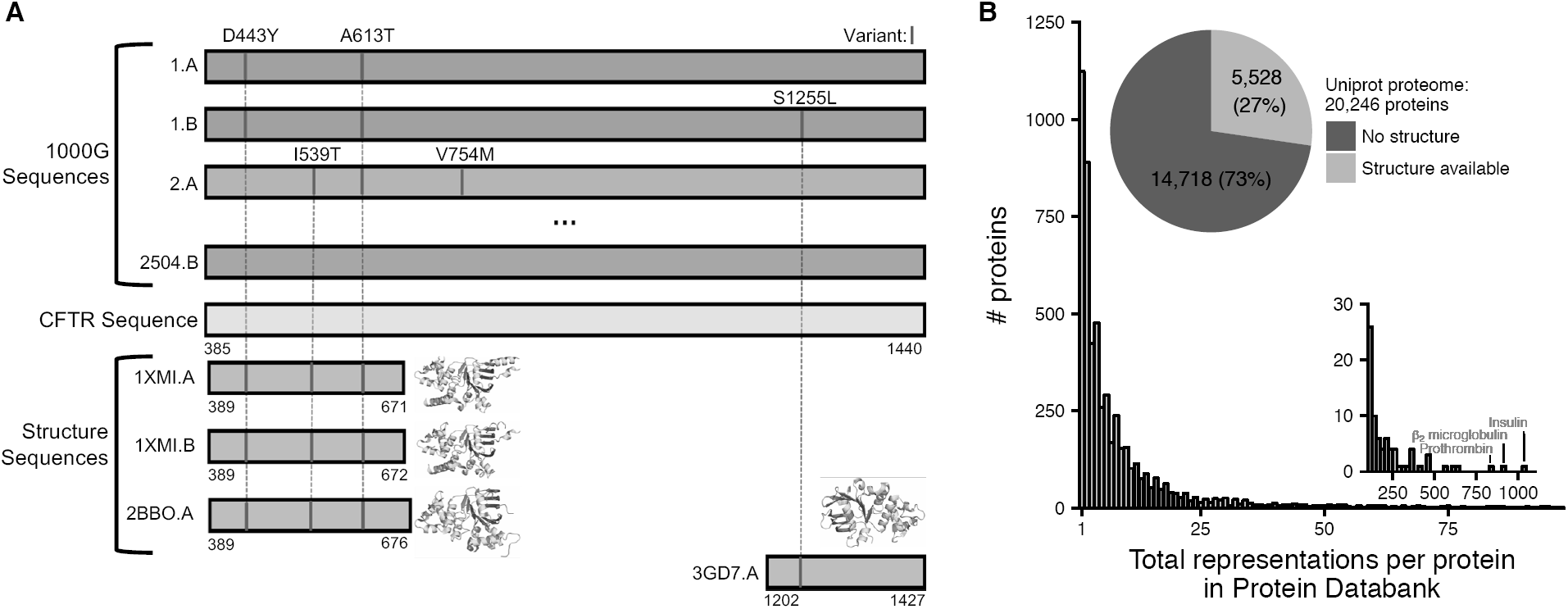
Human protein sequence coverage by structures. (A) Schematic of the mapping of human protein sequence variants observed in the 1000 Genomes Project (1000G) onto sequences represented by different structures in the PDB. The 1000G sequences come from 2504 humans from diverse populations. Here, the cystic fibrosis transmembrane regulator protein (CFTR) is used to illustrate the process. Our analyses focus on regions aligned between protein sequences and residues with resolved coordinates in structures. Some protein variants (e.g., CFTR V754M) are not covered by structures while others (A613T) are covered by multiple structures. We obtained similar results considering all structures and a non-redundant set of structures. (B) Coverage of the human proteome by structures (pie chart). For proteins with structural coverage, the histogram gives distribution of the total number of structures available for each protein. Extreme examples with greater than 100 structures available are presented in the inset.

### Available structures do not represent all protein sequences found in human populations

To quantify how well available protein structures represent human individuals, we computed the percent of observed amino acid sequences in 1000G individuals that match the sequence in each available structure. On average, over the 66,971 structures considered, the sequence modeled in the structure represents 74% of the sequences observed in 1000G.

The sequences of 15,540 structures (23%) do not match the sequences of *any* individual in 1000G (Figure 2A; Figure S3A). Mutations are often engineered into sequences to aid in crystallography or to reflect a variant of interest; 26% of human protein structures (17,524 out of 66,971) contain annotated engineered mutations. After masking engineered mutations, 2,691 structures (>4%) still do not match the sequence of any individual, and less than 15% of structures represent all sequences present in 1000G individuals (Figure 2A). The fraction of human protein structures deposited in the PDB each year that do not match the sequence of anyone in 1000G has decreased after 2005 from ~10% to less than 5% (Figure S4).

**Figure 2.**
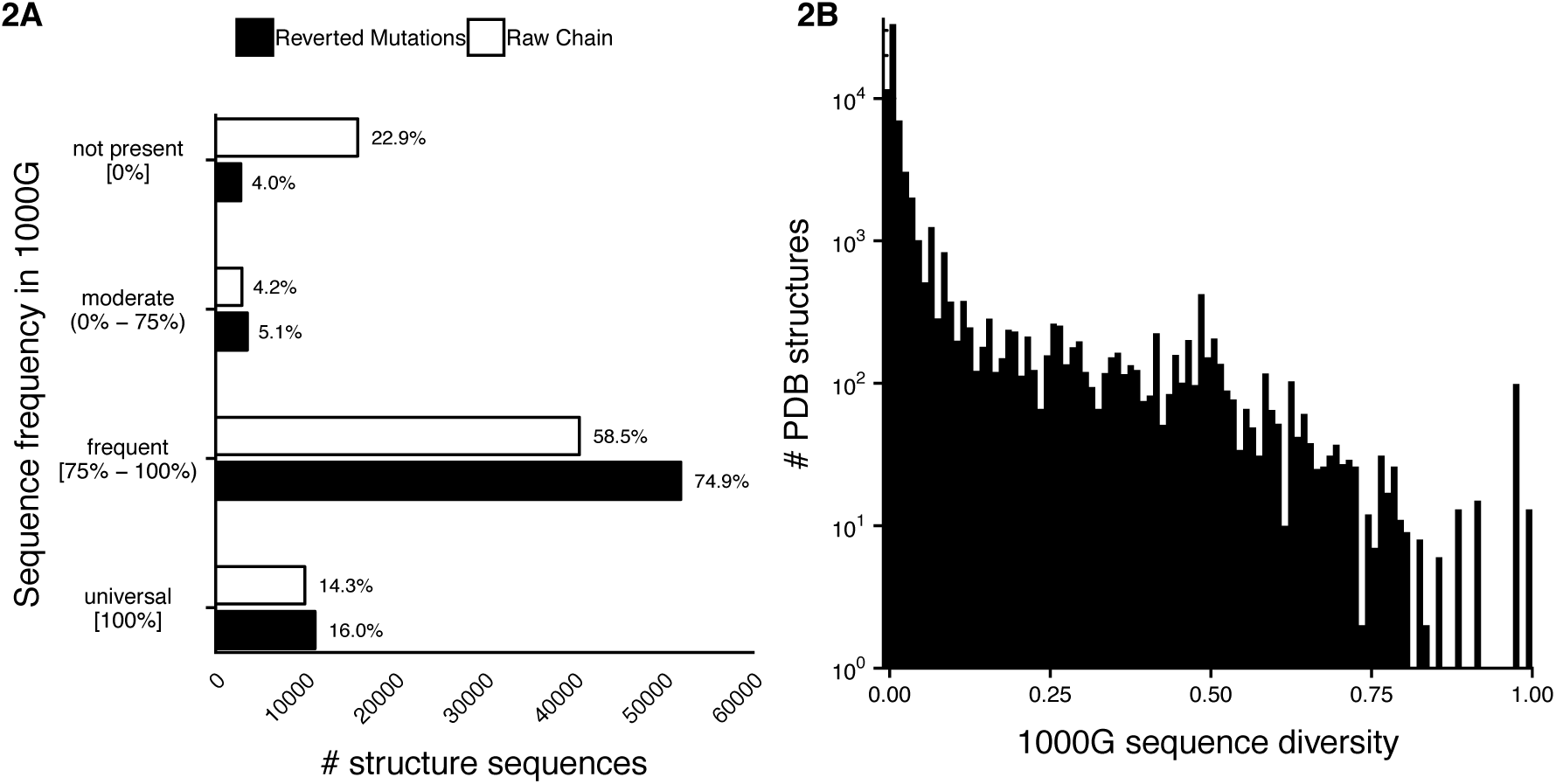
The frequency of protein sequences represented in available structures across diverse individuals. (A) Histograms of the frequency across individuals in 1000G of each sequence represented in a human protein structure. More than 15,000 structures represent sequences not present in any 1000G individual, and only ~15% of structures represent all individuals in 1000G. The white bars reflect the frequencies of the raw sequence modeled in the protein structure across 1000G individuals. The black bars show the 1000G sequence frequencies after masking engineered mutations in the protein structure. The structures are binned based on their sequence’s frequency across 1000G. (B) Histogram of sequence haplotype diversity computed across 1000G individuals for sequences covered by human PDB structures. Higher values indicate greater sequence diversity. Thousands of proteins represented by structures have high levels of diversity across human populations.

To identify structures representing proteins with the greatest underlying human sequence diversity and quantify how well structures capture this diversity, we computed two metrics of sequence-level diversity: the multi-locus Fixation Index (F_ST_) and the haplotype diversity (Browning and Weir, 2010). While most proteins have relatively low diversity, thousands of proteins with structures have high sequence diversity across humans (Figure 2B; Figure S3B). Examining the most sequence-diverse structures, the 293 in the top 10% for both F_ST_ and haplotype diversity, reveals a variety of folds, functions, and patterns of variation (Figure 3; Figure S4). These proteins include diverse functions—e.g., 99 involved in responding to stimulus, 55 in development, and 54 in immune system processes—and diverse binding partners—e.g., 108 protein binding, 94 ion binding, 38 nucleotide binding, and 35 drug binding proteins (Figure 3A).

**Figure 3.**
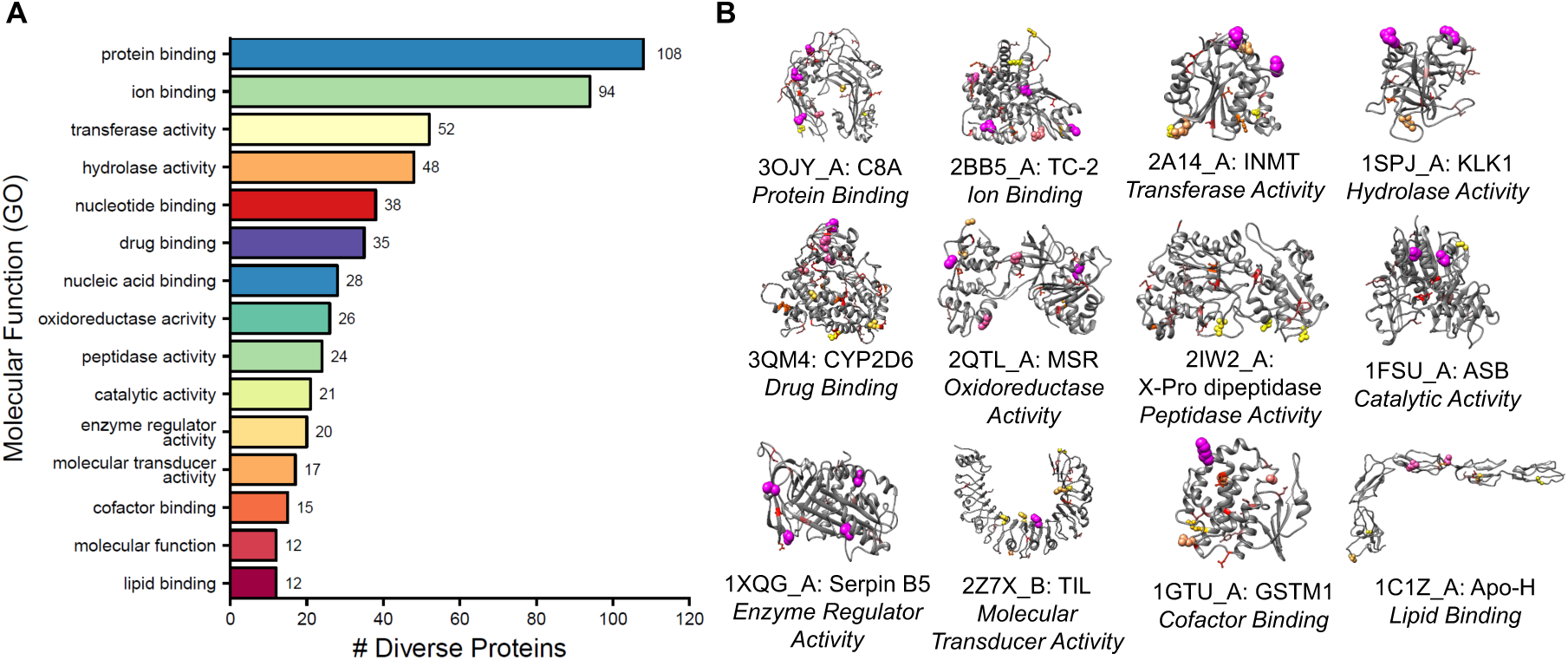
Proteins with highly diverse sequences across human populations cover diverse structural folds and functions. (A) Gene ontology (GO) molecular function slim annotations for structures with greatest diversity across 1000G individuals (top 10% for both F_ST_ and haplotype diversity) highlights a range of functions. (B) Example structures illustrate the diverse folds and variant distributions across functional categories. Color and scaling are graded by allele frequency in 1000G (large purple spheres: ≥ 10% MAF; small yellow spheres: 1% ≤ MAF < 10%; red sticks: 0.1% ≤ MAF < 1%). The PDB ID and protein name are given with the molecular function they illustrate. Additional examples are given in Figure S4.

While we focus on single nucleotide variants here, larger variants are also common in protein-coding sequences. These large variants often dramatically alter the protein structure, removing one or more domains or even leading to complete loss of expression due to nonsense mediated decay (NMD). To approximate the magnitude of differences between protein structure and 1000G sequences, we collected all frameshift, early stop, and in-frame deletions from 1000G that affected one or more residue in a PDB protein structure. Not including variants predicted to cause NMD, our analysis found that 505 unique variants caused multi-residue changes in one or more PDB structures, and 73 of these variants were common. The most frequent consequences were stop gain variants followed by frameshift, and then in-frame deletions.

Thus, it is common for available protein structures to differ from the amino acid sequence present in many humans, even in the relatively small sample from 1000G. However, the degree to which these differences impact structure and function depends on their context and extent. Therefore, in the next sections, we evaluate the structural and functional relevance of these sequence differences. We focus on the “reverted” sequences in which engineered mutations are reverted to original annotated residue. However, users of protein structures often do not account for these engineered mutations, and thus where appropriate, we also present results for “raw” sequences taken directly from structures.

### African ancestry protein sequences are less likely to be represented by available structures

Given that protein sequence variation found in human populations is rarely fully captured by available structures, we explored whether sequences observed in different human populations differ in their likelihood of being represented in structures. Individuals with European ancestry are overrepresented in most genomic databases, and African ancestry individuals are often the least represented (Popejoy and Fullerton, 2016). Thus, we hypothesized that protein sequences from individuals of European ancestry would be more likely to be represented by current 3D structures than those from other populations, especially those with African ancestry. To test this, we computed the fraction of individuals represented by each reverted PDB structure within each of the five super-populations in 1000G (AFR: African, AMR: Admixed American, EAS: East Asian, EUR: European, SAS: South Asian).

All populations have a significant number of protein sequences that are not represented by each structure; however, protein structures are significantly less likely to represent versions of a sequence observed in African ancestry populations than in other human populations. For example, the fraction of structures matching all AFR sequences is 34% compared to 38% for EUR (Figure 4A). Differences were similar when comparing to all other populations (Figure S5A, *P* < 4.5 × 10^−16^ for AFR vs. each other population, Fisher’s exact test with Holm correction). This bias is also present among the non-redundant structures and the raw sequences (Figure S5A,B). Overall, there was also a modest, but significant difference in the fraction of individuals matched on average by a structure between AFR and non-admixed Eurasian populations (raw: 73.5% vs. 74.0%, reverted: 91.5% vs. 92.1%).

**Figure 4.**
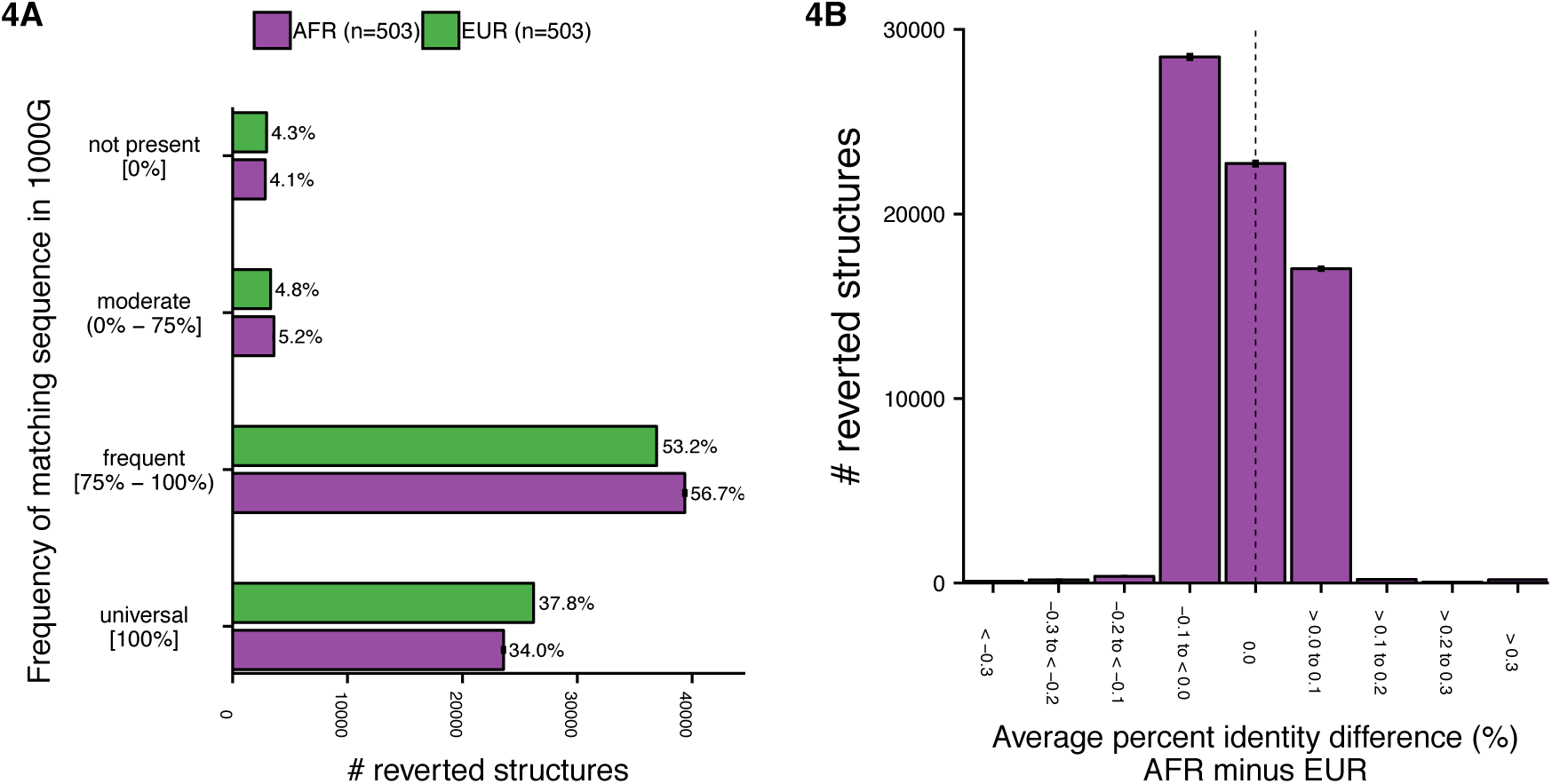
Available protein structures are less representative of sequences from individuals of African ancestry than sequences from individuals from other populations. (A) Histogram of the frequency of protein sequences represented by available structures in African (AFR) and European (EUR) populations after reverting engineered mutations. African individuals were down-sampled to match the smaller European population (503), and the average over ten random samples is presented with standard errors. Available protein structures are significantly less likely to represent sequences of African individuals than of European individuals (*P* < 4.5 × 10^−16^, Fisher’s exact test with Holm correction). Individuals of African ancestry are also significantly under-represented compared to each other population (Figure S5A,B). (B) Histogram of the difference in average percent sequence identity to the sequence represented in available structures between African and European individuals. The protein sequences in African ancestry individuals are modestly, but significantly less similar to sequences modeled in 3D structures than sequences from European-ancestry individuals (*P* < 9.9 × 10^−324^, binomial test) and all other populations (*P* < 9.9 × 10^−324^ for all, Figure S5C).

Additionally, Africans have significantly lower average percent sequence identity to available structures than individuals from other populations (*P* < 2.2 × 10^−16^ for each, Wilcoxon signed-rank test), and African sequences are 1.65 times more likely to have a larger sequence difference compared to other populations (AFR vs. EUR in Figure 4B; all pairs in Figure S5C; *P* < 9.9 × 10^−324^ for all, binomial test). This large consistent difference is not seen in comparisons for any the other populations (Figure S5C).

Surprisingly, Europeans were only modestly more likely to be represented by available structures than other non-African populations (Figure S5). Thus, in contrast to biases in many genomic databases, the biases in protein structural representation are more uniform across populations, with slightly higher levels in African ancestry individuals, likely due to the greater genetic diversity of African populations.

While differences between individuals’ sequences and available structures are common, the overall sequence differences are often small. The average difference between a structure’s sequence and sequences observed across individuals is 1.1 positions after reverting engineered mutations and 1.3 with unmodified raw structure sequences (Figure S6). In the following sections, we explore how often these relatively small sequence differences are likely to influence structure and function.

### Protein variants that are not represented in available structures influence protein function

Thousands of protein-coding variants that cause Mendelian diseases are known to disrupt structure; in contrast, we investigate how often variants found in individuals without severe disease and not modeled in available structures are likely to influence structure and function (Table 2; Figure 5; Figures S7-9).

**Table 2.** Thousands of common protein variants not represented in protein structures are likely to influence structure and function. Functional residues come from annotations in Uniprot or from the structure itself. The total number of functional residues affected by 1000G variants across all models in the PDB (including redundancy), unique protein positions (no redundancy), and unique proteins (out of 5,528 with structures) are given. Phenome-wide association studies (PheWAS) over the UK Biobank have identified 89 phenotype associations for these specific variants and for 628 variants in these proteins.

| Function | # Structure Positions | # Protein Positions | # Proteins | # PheWAS Associations |
| --- | --- | --- | --- | --- |
| Phosphorylation | 1,631 | 94 | 89 | 1 |
| Disulfide Bond | 1,528 | 134 | 113 | 2 |
| DNA Binding | 2,627 | 322 | 148 | 2 |
| Ligand Binding | 9,105 | 1,463 | 805 | 12 |
| Protein Binding | 49,927 | 6,144 | 1,836 | 72 |
| **Total** | **62,676** | **7,713** | **2,359** | **89** |

**Figure 5.**
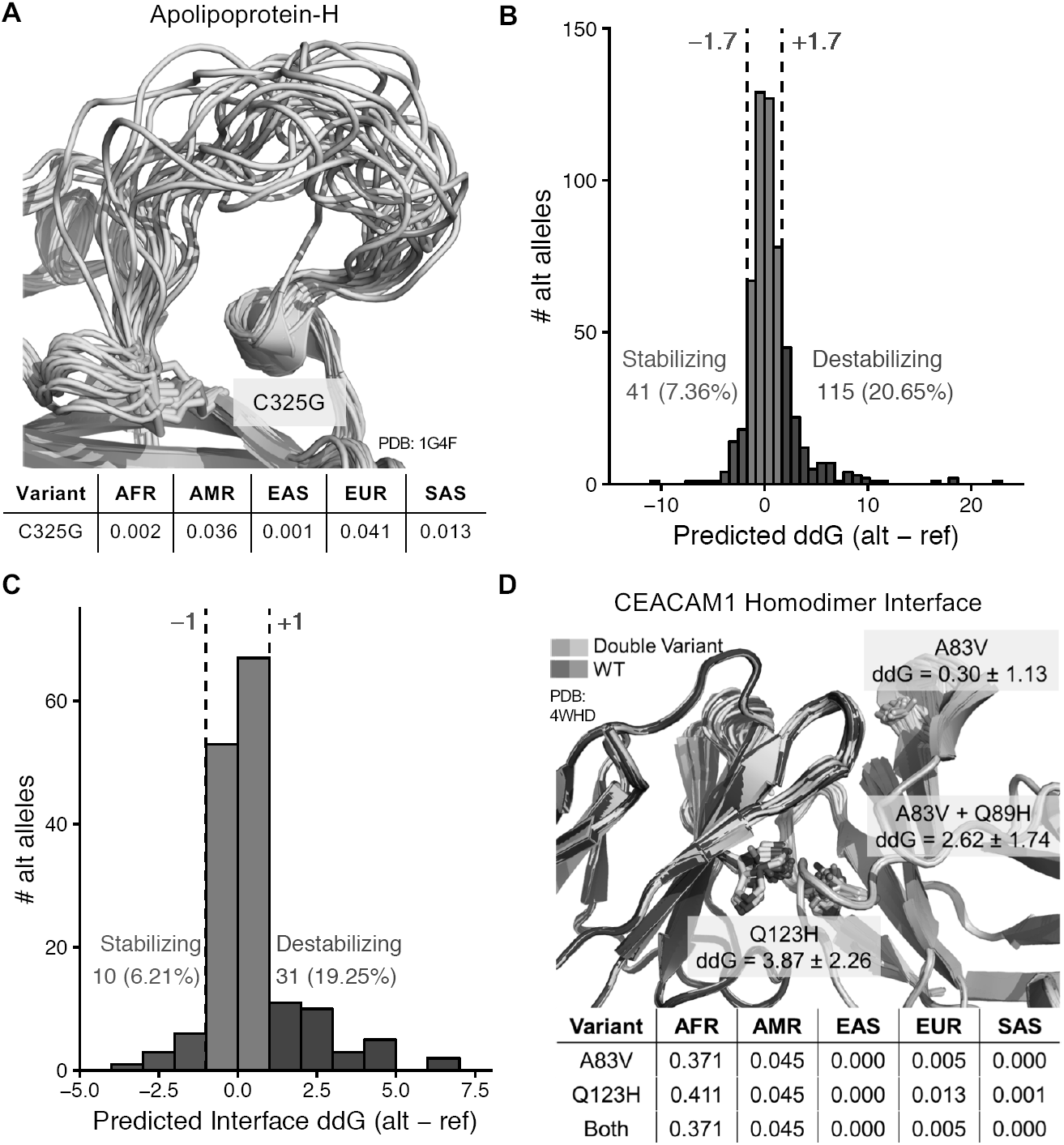
Many unmodeled variants influence protein structure and function. (A) An unmodeled variant (green) removes a disulfide bond in the protein Apolipoprotein-H (APOH; PDB: 1G4F). All conformations in the NMR ensemble are superimposed to highlight variability of the neighboring loop. (B) Histogram of computationally predicted changes in fold stability (ΔΔG) over 557 missense variants in monomeric structures of high quality (<2.5Å) exhibiting higher than average diversity across 1000G. Variants are assigned categories (stabilizing, benign, destabilizing) with an absolute threshold of 1.7 Rosetta Energy Units (REU) and colored according to this cutoff (Methods; Figure S10). We note that a few extreme ΔΔGs, including one >50 REU not pictured, are not physiologically realistic, but likely due to reduced accuracy for highly destabilizing mutations. (C) Histogram of predicted protein-protein interface stability change (Interface ΔΔG) for 161 1000G missense variants within protein-protein interfaces. Variants are assigned to the stabilizing and destabilizing category according to an absolute cutoff of 1.0 REU (Methods). (D) Two co-occurring variants (A83V and Q123H) within the homodimer binding site of CEACAM1 that are common in Africans, but rare in European- and Asian-ancestry individuals are predicted to destabilize the interaction. Predicted binding energy changes for each individual variant and both variants are shown in REU along with standard deviations across all models. All models generated by the flex ddG protocol are superimposed. The frequencies of the variant(s) in 1000G super-populations are given below each structure in (A) and (D).

To illustrate this scenario, we first present an example before moving on to large-scale analyses. Apolipoprotein-H (APOH) is a multifunctional lipid binding protein that is involved in many physiologic pathways including lipoprotein metabolism, coagulation, and the production of antiphospholipid autoantibodies. APOH has substantial variation in its coding sequence, and these variants show significantly different frequencies between human populations. For example, the variant C325G is common (>4%) in European populations, but very rare in both Africa and Asia. This variant is not modeled in any available 3D structures, but it has been shown to disrupt phosphatidylserine binding (Sanghera et al., 1997). We found that it disrupts a disulfide bond tethered to one end of a loop that shows considerable conformational flexibility in an NMR structure (Figure 5A). Furthermore, this variant is associated in several genome-wide association studies (GWAS) with blood protein and cardiometabolic traits, including plateletcrit and LDL cholesterol levels (Bycroft et al., 2017; Consortium et al., 2013; Iotchkova et al., 2016; Suhre et al., 2017). These associations demonstrate the structural and physiological relevance of this unmodeled variation.

We also identified many high frequency variants that have previously been crystalized and shown to significantly impact protein function and downstream analyses, but are not commonly considered. For example, Glutathione S-transferase P (GSTP1) is a cytosolic enzyme that protects cells from xenobiotic substances that has two common variants (I105V and A114V). Studies of ethacrynic acid (EA), an inhibitor of GSTP1 that failed to pass phase II clinical trials due to severe side effects, and phenethyl isothiocyanate (PEITC), an anti-cancer drug with a short half-life due to efficient metabolism by GSTA1, have been examined in complex with Ile105 and not Val105 (Ang et al., 2009; Kumari et al., 2016). Similarly, we previously demonstrated that several available structural models and analyses of Galectin-8 (LGALS8), a beta-galactoside-binding protein involved in cellular defense, consider sequences that do not ever occur in human populations (Ely et al., 2018).

Failure to account for common variants can and does influence conclusions based on available structures and, given their differences in frequency across populations, bias conclusions to specific individuals. As we demonstrate in the next section, such effects on structure and function are common.

### Thousands of protein variants that are not represented in available structures likely influence protein function

We quantified the functional effects of unrepresented variants on a larger scale using four complementary approaches: 1) overlap with functional annotations, 2) phenome-wide association studies (PheWAS) on unrepresented variants using the UK Biobank, 3) variant effect prediction algorithms, and 4) computational modeling of variant effects on protein stability and binding.

First, we examined five annotation types that cover different aspects of protein function that may be influenced by genetic variation: DNA binding sites (Barrera et al., 2016), ligand binding sites (Kang et al., 2015; Magliery and Regan, 2005), protein interaction interfaces (Kann, 2007; Scott et al., 2016), disulfide bonds (Betz, 1993; Li et al., 2013), and phosphorylation sites (Cohen, 2001; Humphrey et al., 2015) (Table 2, Methods). We identified 322 missense 1000G variants at positions that influence DNA binding sites across 148 proteins. Due to the redundancy in the PDB, this includes a total of 2,627 positions in structures that modify DNA binding sites. Similarly, analyzing small ligand binding sites, we found 9,105 1000G variants representing 1,463 unique protein residues across 805 proteins in annotated ligand binding sites. Among these, we observed a substantial degree of functionally relevant polymorphism in CYP enzymes with structural coverage (35 distinct variants near ligands in 14 proteins) and drug binding sites (362 drug binding positions across 217 proteins). We further identified 49,927 variants, covering 6,144 unique positions across 1,836 proteins that influence protein-protein interaction (PPI) interfaces. We identified 134 unique variable residue positions across 113 proteins that mutate a cysteine participating in an annotated disulfide bond. Finally, we identified variants at 1,631 known phosphorylation sites in structures, representing 94 unique positions across 89 proteins which changed a serine, tyrosine, or threonine to a residue that could not be phosphorylated. See Figure 5A and Figure S7 for examples.

Next, we intersected all 1000G missense variants considered in our analyses with the results of a phenome-wide association study (PheWAS) performed over hundreds of traits in 408,455 white British individuals from the UK Biobank cohort. Over all 1000G variants with frequency 0.001 or greater that mapped to PDB structures, 628 have one or more associations, 436 of which are common in 1000G. Intersecting these 628 variants with our collection of sites of interest, we found 89 overlaps: 72 were within a protein-protein interface, 12 near a ligand, two near a nucleotide, and two disrupting disulfide bonds (Table 2). Additionally, 16 were predicted to change protein stability (see below). Given that PheWASs do not necessarily detect the causal variant, but often variants in high linkage disequilibrium with the causal variant, we suspect that the functionally annotated variants likely explain more than 89 of the associations. For 59 of these variants, both reference and alternate alleles are present in different PDB structures, suggesting that some effort has been made to examine structural differences.

We also applied nine variant effect prediction algorithms to evaluate the potential functional effects of alleles not represented in available structures beyond existing knowledge of functional sites (Methods). Overall, 79% of 1000G variants not represented in structures (42,853 / 54,408) are predicted to influence function by at least one algorithm, and 27% (14,516 / 54,408) of these variants are predicted to be functional by more than half of the algorithms (Figure S8). When limiting the analysis to variants with at least 1% frequency in any population, 63% are predicted to be deleterious by at least one classifier (3,128 / 4,982) and 12% (619 / 4,982) by more than half. However, we note that these algorithms have been shown to have only modest accuracy and considerable disagreement (Dong et al., 2015; Tang and Thomas, 2016).

### Computational modeling predicts that unrepresented common variants often influence folding and protein-protein interaction stability

Missense variants commonly affect protein stability, and altered stability often underlies the functional consequences of single residue variants (Martelli et al., 2016; Teilum et al., 2011). To obtain estimates of the effects of unrepresented alleles that are not based on existing annotations, we computationally assessed whether common unrepresented variants are likely to alter the stability of protein 3D structures.

We computed the change in free energy of protein folding between the two alleles (ΔΔ*G*) for 557 variants with higher than average sequence diversity using the Rosetta modeling suite (Kellogg et al., 2011) (Methods). To focus on structures in which ΔΔ*G* estimates are most accurate, we analyzed variants in monomeric structures of high quality (<2.5 Å). Of the 557 variants, 28% (156) significantly altered the stability of the protein in our models; 115 (21%) were destabilizing (positive ΔΔ*G*) and 41 (7%) were stabilizing (negative ΔΔ*G*) (Figure 5B). Figure S9 illustrates these results with three representative examples: one predicted to be stabilizing and two predicted to be strongly destabilizing. We determined the thresholds for defining mutations as stabilizing or destabilizing based on four curated datasets of experimentally determined single point mutation ΔΔ*G*s (Figure S10). We report ΔΔ*G* in REU rather than transforming to kCal/mol to emphasize that our results are based on computational modeling.

In addition to folding stability, we modeled the effects of variants with high diversity across populations on the stability of PPIs. We used the flex ddG method in the Rosetta modeling suite (Barlow et al., 2018) to predict changes in protein-protein binding affinity for 161 interface variants (Methods). Of these variants, 31 (19%) were predicted to destabilize the protein-protein interaction and 10 (6%) to stabilize the interaction (Figure 5C).

For example, we modeled the effects of two commonly co-occurring variants (A83V and Q89H) within the homodimer binding site of carcinoembryonic antigen-related cell adhesion molecule 1 (CEACAM1). CEACAM1 regulates cell adhesion and signaling in a range of contexts via adopting different oligomeric states. The regulation and formation of these conformations is incompletely understood (Bonsor et al., 2015). The two variants are not represented in the structure, but are very common in African populations (>37%) and extremely rare in Eurasian populations (<1%, Figure 5D). The Q89H variant alone and in combination with A83V were both predicted to significantly destabilize the dimer interaction. Thus, common variation likely influences transitions between oligomeric states differently in different human populations.

The analyses presented in this and the previous section indicate that there are thousands of missense variants in human populations that do not cause severe disease, but are likely to affect the results of structure-based studies that only consider structures from single “reference” sequences. These results highlight the need to increase appreciation and communication of the diversity of protein sequence and structure and its potential to influence function. To facilitate the evaluation of whether a structure represents sequences present in a population of interest, we provide the frequency of each PDB structure’s sequence in each 1000G population (Table S1).

## DISCUSSION

Protein structures are critical to experimental and computational analyses of the mechanisms of protein function and to the development of new drugs and therapeutic approaches. However, there are many recognized limitations of the static, incomplete, and non-physiological snapshot of proteins provided by available structural models (Acharya and Lloyd, 2005; Davis et al., 2008; Mannige, 2014). Here, we identify a further challenge: many human protein structures do not represent the diversity of sequences present in the human population, and thus failure to consider human genetic diversity in structural analyses can propagate biases against unrepresented individuals. Furthermore, we find that available protein structures are modestly, but significantly, less likely to represent sequences found in individuals of African ancestry than in all other populations considered. This suggests that, unlike many other databases with biases caused by an oversampling of European ancestry individuals, the differences for structures are driven by the greater genetic diversity of African populations rather than other systematic or sampling biases. While the biases in genomic studies of disease have garnered much attention, our results illustrate the potential for bias and the challenges to reproducibility in basic science investigations that rely on other “reference” databases.

Many rare protein variants influence protein structure in ways that lead to disease, and variants occurring in databases like 1000G have often been used to represent “benign” mutations. Our results demonstrate that thousands of variants common in human populations are likely to influence diverse aspects of protein fold, stability, and interactions. These variants may not always influence disease risk, but they are often physiologically relevant. For example, otherwise benign variants in cytochrome P450 (CYP) enzymes have critical effects when individuals carrying them exposed to exogenous compounds, necessitating a personalized dosing approach to address the different pharmacokinetic properties across individuals caused by single residue changes (Preissner et al., 2013; Zanger and Schwab, 2013). Structural analyses often focus on precise characterization of the functional activity of a single protein or complex. Such *in vitro* and *in silico* analyses that rely on protein structures drawn from a single “wild type” sequence may be sensitive to small sequence differences that can substantially impact conclusions. Thus, just as we often consider proteins as conformational ensembles rather than single static structures, we also must consider protein “sequence ensembles,” each with their own structural and functional properties. The potential impact of the biases we identify is substantial; the 293 most diverse human protein structures have been cited by over 40,000 papers in PMC.

Our results underestimate the functional effects of human genetic variation on modeled protein sequences for several reasons. First, we limit our analyses to missense variants. Other variants with potential to cause profound impacts on protein structure (Cooper et al., 2010), such as insertions, deletions, early stop codons, and copy number variants, are common throughout the 1000G and other exome sequencing data sets (1000 Genomes Project Consortium et al., 2015; Lek et al., 2016; Ruderfer et al., 2016). Furthermore, in some cases, synonymous variants can impact protein structure and function through changes in mRNA stability and dynamics (Sauna and Kimchi-Sarfaty, 2011). We also have not quantified the effects of variants that influence splicing. Alternative splicing creates significant additional structural and function diversity in the human proteome, and variants that influence splicing often contribute to disease (Li et al., 2016; Ongen and Dermitzakis, 2015; Park et al., 2018). Finally, while the 1000G cohort contains individuals from diverse populations, it does not fully capture the diversity of human populations, especially within Africa and South Asia (Crawford et al., 2017; Mallick et al., 2016). Deeper sequencing of already studied populations and the sequencing of additional populations from around the globe will further increase our knowledge of structurally relevant genetic variation.

We are unlikely to be able to experimentally model each individual’s unique proteome in the near future. A diverse array of algorithms for predicting the effects of protein-coding mutations have been developed over the past several years (Figure S8); however, these methods often disagree and likely perpetuate biases found within the datasets on which they were trained (Cline and Karchin, 2010; Mahmood et al., 2017; McCarthy et al., 2014). Furthermore, most provide only qualitative predictions about variant effects that do not enable additional mechanistic or structural analyses. Our results suggest that combining computational methods for modeling the effects of sequence changes with representative structures is a promising approach for addressing this challenge. Furthermore, investigators must consider genetic differences and the potential for a protein structure to reflect only a subset of the human population when sequences are selected for experimental characterization to avoid propagating health disparities. We propose that PDB structures should include an additional annotation that quantifies human variability within the structure and its sequence’s prevalence across populations.

While it is convenient to model proteins as single, canonical amino acid sequences, in reality this is often inaccurate and presents a roadblock to achieving the goals of both basic protein science and personalized medicine. The costs of not accounting for human variation in proteins that significantly impact drug efficacy and toxicity are becoming increasingly appreciated (Chan et al., 2016; Perera et al., 2014; Petrovski and Goldstein, 2016; Roden, 2016; Tekola-Ayele et al., 2015). In these cases, the effects are often profound and readily apparent in a system-wide response. However, our results suggest that subtle effects are common in diverse proteins, and that they are likely to influence many experimental assays and *in silico* analyses of protein function. Accounting for human genetic variation early in the research and development pipeline can have significant cost saving and safety benefits (Hauser et al., 2018). In short, human genetic variation must be explicitly accounted for in structural studies to ensure results that accurately describe protein structure and function across all human populations.

## METHODS

### Mapping variants to protein structures in the human proteome

We used Uniprot’s annotated human proteome (UP000005640_9606, https://www.uniprot.org/help/human_proteome) as the human proteome for all analyses. At the time of our download (February 22, 2018), it contained 20,246 human proteins.

Several tools are available that enable mapping known non-synonymous single nucleotide variants onto PDB structures and functional annotations (Prlic et al., 2016; Ryan et al., 2009; Sivley et al., 2018). To map genetic variants from 1000G (1000 Genomes Project Consortium et al., 2015) into available protein structures, we used the previously described PDBMap pipeline (Sivley et al., 2018). In brief, variant consequences were predicted using v82 of Ensembl Variant Effect Predictor (GRCh37.24) (McLaren et al., 2016) and all missense variants across the proteome were collected. Variant positions represented within the corresponding Ensembl transcript model were mapped onto PDB structures using the reference sequence to PDB sequence alignment provided by SIFTS (Structure Integration with Function, Taxonomy and Sequence) (Velankar et al., 2013). Discrepancies between the transcript model and these alignments were resolved with Needleman-Wunsch optimal pairwise alignment. In cases where a single residue had several alternative amino acids, a single alternative amino acid was randomly selected. For all variants, sample-specific alleles were determined for all 5008 versions of each protein across the 2504 individuals in the 1000G phase 3 dataset. For raw chain sequence analyses, these individual-based sequences were compared directly with the PDB chain sequence defined by all residues with at least one atom with defined coordinates. Any residues lacking coordinates in the PDB structure (due to gaps, etc.) were not considered, even if they appeared in the SEQRES in the structure. Protein structural models often do not cover the entire protein sequence due to experimental limitations or specific interest in a particular protein domain rather than the entire protein. In our analyses, the average coverage of a protein sequence by a structural model was 54.2%. In general, smaller proteins (<500 amino acids) had greater sequence coverage than larger proteins (Figure S1).

To resolve ambiguities in the raw structures and sequences in the PDB, we used the following protocol:

1. Structures containing multiple biological assemblies or models were represented by the first listed model of their asymmetric unit for all sequence-based analyses. For functional annotations based on residue pair distances, all physiologically relevant biological assemblies and all models were considered.
2. Atoms appearing multiple times within the same residue due to alternate positions were represented by the first set of coordinates only.
3. Residues appearing multiple times within the same chain due to insertion codes were represented once for all sequence-based analysis. If one of the insertion-code amino acids matched the transcript sequence, this amino acid was selected for this position; otherwise, the amino acid with the lowest insertion code was selected. For all functional annotations based on residue-pair distances and energetic calculations, all insertion code residues were included.
4. For energetic calculations, residues lacking one or more backbone atoms were removed.
5. Non-canonical amino acids were represented as their canonical counterpart when possible.
6. Engineered missense mutations were reverted to their listed original amino acid for all sequence-based analyses using “reverted” sequences, regardless of whether the original amino acid matched the transcript or not. Engineered insertions/deletions were not altered or considered. Missense mutations were identified through the PDB’s REST interface under the “mutation” tag. Engineered mutations may be introduced for a variety of reason. For example, the common N700S polymorphism in Thrombospondin 1 (THBS1) has been linked to increased risk of premature coronary artery disease and was introduced to the structure 2RHP specifically to examine structural differences resulting from the variant (Carlson et al., 2008). The structure of FBS1 (PDBID: 5B4N chain A), a component of the SCF ubiquitin ligase, contains 40 engineered mutations. These variants are found in FBG3 and were introduced into FBS1 to study structural and functional differences in the loop regions of these homologous proteins (Nishio et al., 2016) and DUSP5 (PDBID: 2G6Z), dual specificity protein phosphatase 5, contains a C263S mutation required for crystallization (Jeong et al., 2007). Additionally, there are cases where engineered mutations are introduced to reflect a variant that has a higher frequency than the reference allele. For example, the variant L47F was introduced into 5BUC to reflect the more common allele (REMARK 999 in PDB header). In total, 17,524 out of 66,971 (26.2%) structures contain one or more annotated engineered missense single residue mutation.
7. All non-human residues identified through the DBREF header information were filtered.

Considering computationally derived models of human proteins based on resolved structures of homologous proteins would increase coverage of the human proteome to >70% (Sivley et al., 2018; Somody et al., 2017). However, because homology modeling requires introducing amino acid changes and the results are of variable accuracy, we limit our structural analysis to experimentally determined human structures from the PDB.

Sequence identities were calculated for all sample proteins individually based on the alignment between the individual transcript accounting for 1000G alleles and the PDB chain sequence accounting for all considerations listed above.

### Non-redundant chain set selection

All PDB structural models were grouped by their corresponding Uniprot ID and sorted by the total number of transcript residues they covered (not counting gaps or SEQRES entries that lacked coordinates for any atoms). The structure covering the most residues was retained. For the remaining set of structures, a structure was retained if it overlapped a previously selected structure by less than 50%. This process was repeated for all Uniprot ID groups. A schematic of this method is presented in Figure S2.

In the current structural knowledgebase, domains or sections of a single protein are sometimes split into multiple structures. For example, insulin’s B chain is often found as a separate structure from the rest of the protein, leading to its high structure count. When generating the non-redundant set, we found 1,146 proteins with coverage split across structures. The most extreme example, Zinc finger protein 268, was covered by 23 separate structures.

### Population analyses

All sequence identity measures were calculated according to the sequence identity of samples in 1000G with the raw or reverted structure sequence. Gaps, residues lacking any atom coordinates, and attached tags that did not align to the corresponding transcript sequence were not considered. Population-level statistics were computed for the five 1000G super-populations: AFR, AMR, EAS, EUR, SAS.

For comparisons of percent identity, we downsampled to the size of the AMR population, the smallest in 1000G, to account for potential sample size artifacts. For this process, a random sample of individuals was taken from each population equal to the size of the AMR (347, both chromosomes per individual). All chromosome pairs were retained in the downsampling. Within each subset, the percent of samples in the population subset that matched the structure sequence exactly was counted. This process was repeated five times and the results presented are the average across all five subsets.

To examine the differences in average sequence identity between populations, 1000G sequences were grouped by population and the average sequence identity within each population was calculated for each PDB chain. Then, average percent identities were subtracted for each chain, yielding a histogram of differences.

### Haplotype diversity and F_ST_ calculations

#### Haplotype diversity

We used the haplotype diversity measure suggested by Masatoshi Nei and Fumio Tajima (Nei and Tajima, 1981):

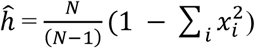

where *N* is the number of individuals in each population, *i* indexes the haplotypes for each chain in each population, and *x*_*i*_ is the corresponding haplotype frequency. We calculated overall haplotype diversity for each chain across all the populations, as well as within each of the 5 super populations defined in the 1000G project.

#### Population F_ST_

We used the Weir & Cockerham method (Weir and Cockerham, 1984) to calculate the pairwise F_ST_ measures for each chain between two populations and an average F_ST_ across all the populations.

We calculated adjusted population size 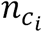 for each population *i* for a sample with *r* different populations using the formula below:

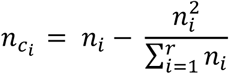

where *n*_*i*_ is the total number of haplotypes in each population.

We then used the adjusted population sample size to calculate an average population size (*n*_*c*_) using the following formula:

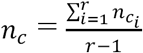

We then calculated weighted average haplotype frequency for each haplotype across all the populations using the following formula:

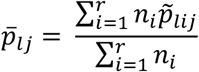

Here, 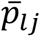 is the average weighted frequency for haplotype *l* in protein chain *j* and 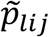 is the haplotype frequency for the same haplotype in each population *i*.

We then used the above parameters to calculate Mean Squares Among (MSA) and Mean Squares Within (MSW) population variation for each haplotype *l* in protein chain *j*, represented by *MSA*_*lj*_ and *MSW*_*lj*_ respectively, as shown in formulae below:

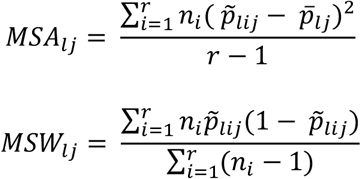

Finally, we calculated the average population F_ST_ for each chain 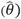 using the following formula:

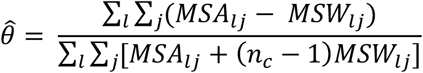

Using the algorithm described above, we calculated population average F_ST_, as well as pairwise F_ST_ where we considered two populations at a time (r = 2).

### Gene ontology annotation of structures with high diversity

Structures with highest diversity were selected by intersecting the top 10% of non-redundant structures ranked by F_ST_ (both average and pairwise) and the top 10% of non-redundant structures by haplotype diversity. This resulted in a high diversity set of 293 proteins from 4,313 structures.

Gene ontology (GO) annotations were assigned for all high diversity structures with the PDB GO annotation set downloaded from the European Bioinformatics Institute (ftp://ftp.ebi.ac.uk/pub/databases/GO/goa/PDB/goa_pdb.gaf.gz, retrieved May 20, 2018). A total of 3,434 structures had attached GO annotations leaving a set of 233 high diversity proteins across 3,434 structures. These GO terms were translated into the Protein Information Resource slim subset (http://www.geneontology.org/ontology/subsets/goslim_pir.obo, retrieved May 20, 2018) with the Map2Slim function of OWLTools (Mungall et al., 2017) and the GO relations defined in the recommended go-basic version (http://snapshot.geneontology.org/ontology/go-basic.obo, retrieved May 20, 2018). The final set of structures for which slim GO annotation counts could be performed was 233 proteins across 3,434 structures. All counts were based on unique GO annotations seen for each protein to avoid structure redundancy.

### Citation Analysis

Citation counts in PMC were generated for the primary citation for each PDB structure as identified directly from the RCSB PDB. First pubmed ID’s were retrieved for each PDB using the REST API. These pubmed ID’s were linked to articles citing them by their PMC ID’s using the Entrez module of Biopython (Cock et al., 2009). Specifically, the E-utility e-link (pubmed_pmc_refs) was used to identify citing articles.

### Functional effect prediction analysis

Variant function effect scores were collected for all structural 1000G missense variants from the dbNSFP database (Liu et al., 2011) through the ANNOVAR platform (Wang et al., 2010). dbNSFP is a database of functional classifications for all potential non-synonymous single-nucleotide variants in the human genome. We used 9 score predictions compiled by dbNSFP: SIFT (Ng and Henikoff, 2001), Polyphen2-HDIV, Polyphen2-HVAR (Adzhubei et al., 2010), LRT (Chun and Fay, 2009), MutationAssessor (Tang and Thomas, 2016), FATHMM (Shihab et al., 2013), MetaSVM, MetaLR (Dong et al., 2015), and PROVEAN (Choi and Chan, 2015).

For the classification counts, we limited our analyses to predictors with an established classification threshold for a functional effect. For classifiers with more than two categories (e.g., Polyphen2 classifies variants as potentially damaging, probably damaging, and benign), we only counted the most extreme category. We excluded MutationTaster from all analyses, because it automatically classifies a variant as benign if it appears as a homozygote in 1000G more than four times.

It is important to note that some variant effect prediction methods have potential drawbacks for analyzing variants in populations such as 1000G. In many cases, they are primarily aimed at predicting system-wide impacts of variants and may downplay subtle effects on protein function that do not lead to a fitness cost. Consequently, variant frequencies in populations such as 1000G may be used as features when training models such as Meta-SVM and Meta-LR. However, given the common use of these methods as a ‘first pass’ filter when analyzing genome-wide datasets and the significant number of variants predicted by multiple methods to be deleterious, they can point to variants with the potential to impact protein function which can further be followed up with more sensitive and high-resolution methods.

### Mapping variants to functional annotations

For all analyses, only chains with 50 or more residues were considered. All Uniprot sequence positions were collected through the SIFTS PDB–Uniprot sequence alignments (Velankar et al., 2013). All Uniprot annotations were parsed from the raw .xml release (ftp://ftp.uniprot.org/pub/databases/uniprot/, retrieved October 23, 2017). Only positions where the chain amino acid matched the sequence amino acid at the corresponding position in the Uniprot database were considered.

#### Phosphorylation Sites

Phosphorylation of serine, tyrosine, or threonine is a common regulatory mechanism (Humphrey et al., 2015), and loss of phosphorylation sites can significantly impact protein function (Cohen, 2001). For example, the variant S513L in medium-chain acyl-CoA synthetase 2 (MACS2) (AFR: 0.13; AMR: 0.14; EUR: 0.07; EAS: 0.20; SAS: 0.16) is associated with risk factors for metabolic syndrome (Lindner et al., 2006). To identify variants influencing potential phosphorylation sites, all variants with a reference amino acid of T, Y, or S and an alternate amino acid other than T, Y, or S were identified. These variants were filtered for those containing Uniprot annotations of “Phosphothreonine”, “Phosphoserine”, or “Phosphotyrosine.”

#### Disulfide bonds

Covalent disulfide bonds between two cysteine residues can provide strongly stabilizing interactions in folded proteins through enthalpic contributions and by altering the entropic landscape (Betz, 1993). In addition to their role in stability, disulfide bonds contribute to protein function and disease through modifying conformational dynamics, e.g., by “locking” regions into functional conformations or tethering domains (Khoo and Norton, 2011; Li et al., 2013). To identify variants influencing disulfide bonds, all variants altering a cysteine at a position with a disulfide bond annotation in the Uniprot database were identified.

#### DNA/RNA binding sites

We identified all residues with atoms within 6 angstroms of DNA or RNA in an available structure.

#### Ligand Binding Sites

Biological assemblies for all protein structures were examined for the presence of ligands and all residues with any atom within 6 angstroms of a ligand atom were identified. The propensity for single residue changes within small molecule binding sites to alter binding (Kang et al., 2015; Magliery and Regan, 2005) is well studied and plays important roles in personalized therapies that draw from polymorphisms found within binding sites of metabolic proteins such as cytochrome P450 (CYP) enzyme (Preissner et al., 2013). Because protein structures are often deposited with non-physiologically relevant artifact ligands carried over from crystallographic analysis, several filters were applied to identify physiologically relevant ligands:

1. All ligands with ‘invalid’ annotation in the Binding MOAD dataset were removed (Ahmed et al., 2015).
2. A manually curated set of common artifact ligands, including ligands frequently annotated as invalid in the MOAD dataset, was generated. The filtered common artifact ligands are: 12P, 144, 15P, 1PE, 1PG, 2CV, 2NV, 2PE, ACY, ARS, AZI, B3P, BCN, BCT, BF2, BME, BNG, BO3, BTB, BU1, BU3, C10, C8E, CAC, CAD, CCN, CDL, CM5, CO3, CPS, CXS, CYN, D12, D1D, DIO, DMF, DMS, DMU, DOD, DPO, EDO, EOH, EPE, ETE, FLC, FMT, FUC, GAI, GAL, GLC, GOL, H2S, HED, HEX, HEZ, HGB, HTO, IMD, IOD, IUM, LDA, LFA, LMT, LMU, MAN, MBO, ME2, MOH, MPD, MPG, MPO, MRD, NDG, NET, NH2, NH4, NHE, NI, NO3, OLC, P33, P4C, P6G, PDO, PE4, PE8, PEG, PEO, PG4, PGE, PGO, PGR, PGW, PO3, PO4, POL, SCN, SGM, SO3, TAM, TG1, TRD, TRT, UNL, UNX, UPL, URE, VO4, XPE, XY1, YT3.
3. PDBbind (Liu et al., 2015) provides a dataset that matches all ligand-bound structures with their binding affinity, when available. Any ligand with an associated binding affinity was retained. Finally, all ligands that did not appear in either MOAD or PDBbind were removed using the common artifact ligand set.

In addition to the broad survey of ligand binding sites, we also examined variants in CYP enzymes and those found at known drug binding sites. Of the 24 unique CYP enzymes with structural coverage in our analysis, 14 had variants near ligands. To identify known drugs, we mapped ligands to their corresponding DrugBank (Law et al., 2014; Wishart et al., 2008) annotation, filtering for compounds with no annotation or that fell within the ‘Experimental’ category (the majority of the resulting ligands were categorized as approved or investigational).

#### Protein-protein interface variants

Due to the critical role of protein-protein interaction (PPI) interfaces in protein signaling, folding, disease, and drug design (Kann, 2007; Scott et al., 2016), all biological assemblies were evaluated for residues at protein-protein interfaces. These residues are those with average side chain coordinates within 8 angstroms of the average side chain coordinates of a residue in a different chain. Since many biological assemblies are included that are not physiologically relevant, we took the following steps to filter non-relevant biological assemblies:

1. We used the QSBio dataset of annotated biological assemblies, which uses similarity and conservation to assess the confidence of quaternary structures across biological assemblies with a focus on homomers (Dey et al., 2018). Any complexes annotated in QSBio as “Very low”, “Low”, or “Medium” confidence were filtered.
2. We also used the PiQSi manual curation of biological assemblies (Levy, 2007). Any assemblies annotated as “YES” under Error were filtered.

### UK Biobank PheWAS Analysis

We intersected all missense variants considered in our analyses with the results of a PheWAS performed over hundreds of traits and millions of variants from 408,455 white British individuals from the UK Biobank cohort. We considered all variants and associations present in the Gene Atlas database on June 19, 2018 (Canela-Xandri et al., 2018). We followed the quality control criteria and thresholds suggested by the Gene Atlas with the exception of using a stricter threshold on imputation quality (0.8 vs. 0.3) due to our interest in rare variants as well as common variants. We further focused on associations found for variants with other evidence of structural effects (e.g., annotated functional sites or significant effects on stability or interactions).

### Modeling of variant effects on protein stability

Protein stability calculations were performed using the most accurate high resolution protocol as identified and described by Kellogg et al (table 1, row 16) (Kellogg et al., 2011). This approach approximates the free energy change between the unfolded and folded protein state for both alleles. The difference in free energy change between alleles, therefore, predicts whether a variant is destabilizing (positive ΔΔ*G*, reduction in the change in free energy) or stabilizing (negative ΔΔ*G*, increase in the change in free energy). In brief, this protocol begins by pre-minimizing all input structures (backbone and sidechain) with harmonic distance constraints on all alpha carbon distances within 9 angstroms of the original structure. Next, 50 iterations of rotamer optimization (repacking) followed by three rounds of gradient-based minimization are performed, beginning with a down-weighted repulsive term and increasing it for each round of minimization. Throughout, original distance restraints are preserved to prevent extreme backbone movements. This protocol allows for some backbone movement with a dampened repulsive term to allow efficient sampling of conformational space surrounding the initial conformation. Finally, ΔΔ*G* calculations were made by subtracting the average pose score of the top 3 reference models from the average pose score of the top 3 variant models. For all ΔΔ*G* calculations, expression tags and initiator methionines (as notated in the SEQADV records) were removed and the same protocol used for the benchmark was performed.

To identify an appropriate threshold for categorizing variants as ‘stabilizing’, ‘neutral’, or ‘destabilizing’, we benchmarked the ΔΔ*G* protocol on a collection of four curated datasets of experimentally determined single point mutation ΔΔ*G*’s (https://github.com/Kortemme-Lab/ddg, retrieved April 17, 2017) resulting in a Spearman’s correlation coefficient of 0.64 (n = 1169) between predicted and experimental ΔΔ*G* values (Figure S7). Based on the tradeoff between sensitivity and specificity, we selected a classification cutoff of magnitude 1.7 ΔΔ*G* (Figure S7A) and assumed that experimental ΔΔ*G* magnitudes of 1.0 or greater indicate effective changes in stability (Figure S7B). We report ΔΔ*G* values in REU as opposed converting them to kCal/mol with a linear model, because this transformation would not change any of our conclusions and using REU emphasizes that these values are computational approximations.

To enrich for variants that are underrepresented in certain populations, we focused on unrepresented variants from proteins in the top quartile of at least one measure of population diversity (average F_ST_ ≥ 0.017 or haplotype diversity ≥ 0.025). This included 25,412 chains covering 2410 unique proteins). We further selected all variants with a frequency of greater than or equal to 0.01 in any population where the corresponding amino acid in the 3D structure was the reference amino acid, yielding 5165 unique variants across 48,067 chains. Finally, to focus on variants and structures in which the ΔΔ*G* computation is most accurate, we analyzed monomers between 50 and 500 residues in length with NMR or X-ray crystallography derived structures with a resolution of 2.5 angstroms or better. A single variant with highest frequency was selected for each structure yielding a final set of 557 unique variants across 557 chains.

### Protein binding affinity prediction

A subset of PPI interface variants were selected for energetic evaluation with the flex ddG protocol (Barlow et al., 2018). To enrich for variants with high diversity, only structures with haplotype diversity greater than or equal to 0.025, average F_ST_ greater than or equal to 0.017, or frequency greater than or equal to 0.01 in any population were included. Only NMR and X-ray crystallography structures with resolution less than or equal to 2.5 angstroms were considered. Additionally, in the interest of computational feasibility, only structures including a total of 1000 residues or less with 100% sequence identity to the transcript(s) were considered. Since this set included many repeated protein positions, a single representative structure was selected for each protein position as the structure covering most residues across both chains in complex or the structure with best resolution. This yielded a set of 456 variants over 402 unique structures and 118 unique proteins.

When predicting energetic consequences of variants in protein interfaces, only the two chains involved in the interface were simulated and variants involved in homodimers were mutated in both chains simultaneously.

The flex ddG protocol was used as described by Barlow et al (Barlow et al., 2018) and draws from the “backrub protocol” (Smith and Kortemme, 2008) to locally sample conformational space around the interface mutation, allowing for backbone changes to accommodate larger side chains. This method was selected as it outperformed other methods when benchmarked with a set of 1,240 mutations with known effects on binding affinity across a range of small to large changes. Briefly, the protocol begins with a minimization of the input structure restrained with harmonic pairwise alpha carbon distances within 9 angstroms in the starting structure. This is followed by a series of backrub moves in the mutation neighborhood, generating models at regular intervals. Each model undergoes side chain optimization of the reference and mutant sequences followed by global minimization. Finally, interface ΔΔ*G* is calculated for each model by subtracting the difference between the reference bound and unbound poses from the difference between the mutated bound and unbound poses. The final ΔΔ*G* for each mutation is taken as the average of all model ΔΔ*G*s. To prevent the disulfide score term from dominating ΔΔ*G* calculations with structures involving disulfide bonds across the interface, we reweighted the disulfide score terms to zero during the ΔΔ*G* calculation steps.

### Structure Visualization

All structures illustrating variants at function positions were generated using PyMOL version 1.8.6.0 (The PyMOL Molecular Graphics System Schrödinger, LLC) and the ray tracing feature. All structures illustrating extreme examples (Figure 3) were generated using UCSF Chimera version 1.11.2. Chimera is developed by the Resource for Biocomputing, Visualization, and Informatics at the University of California, San Francisco (supported by NIGMS P41-GM103311) (Pettersen et al., 2004). Atoms were colored and sized according to overall variant frequency across all populations using render by attribute.

## ACKNOWLEDGMENTS

This work was support by an NIH training grant (T15LM007450) to GRS, NIH R35GM127087 to JAC, NIH R01 NS095989 to CRS, a Vanderbilt Ingram Cancer Center Ambassador’s grant to JAC, NIH R01 GM080403 and R01HL122010 to JM, NIH R01GM126249 to WSB, and the NIH Undiagnosed Diseases Network (NIH U01HG007674).

We thank Rocco Moretti, Chris Moth, Souhrid Mukherjee, and Jonathan Sheehan for helpful discussions and manuscript suggestions. This work was conducted in part using the resources of the Advanced Computing Center for Research and Education at Vanderbilt University, Nashville, TN.

## REFERENCES

1000 Genomes Project Consortium, Abecasis GR, Altshuler D, Auton A, Brooks LD, Durbin RM, Gibbs RA, Hurles ME, McVean GA. 2010. A map of human genome variation from population-scale sequencing. Nature 467:1061–73. doi:10.1038/nature09534

1000 Genomes Project Consortium, Auton A, Brooks LD, Durbin RM, Garrison EP, Kang HM, Korbel JO, Marchini JL, McCarthy S, McVean GA, Abecasis GR. 2015. A global reference for human genetic variation. Nature 526:68–74. doi:10.1038/nature15393

Acharya KR, Lloyd MD. 2005. The advantages and limitations of protein crystal structures. Trends Pharmacol Sci 26:10–4. doi:10.1016/j.tips.2004.10.011

Adzhubei IA, Schmidt S, Peshkin L, Ramensky VE, Gerasimova A, Bork P, Kondrashov AS, Sunyaev SR. 2010. A method and server for predicting damaging missense mutations. Nat Methods 7:248–9. doi:10.1038/nmeth0410-248

Ahmed A, Smith RD, Clark JJ, Dunbar JB, Carlson HA. 2015. Recent improvements to Binding MOAD: a resource for protein-ligand binding affinities and structures. Nucleic Acids Res 43:D465–9. doi:10.1093/nar/gku1088

Ang WH, Parker LJ, De Luca A, Juillerat-Jeanneret L, Morton CJ, Lo Bello M, Parker MW, Dyson PJ. 2009. Rational design of an organometallic glutathione transferase inhibitor. Angew Chemie Int Ed 48:3854–3857.

Apweiler R, Bairoch A, Wu CH, Barker WC, Boeckmann B, Ferro S, Gasteiger E, Huang H, Lopez R, Magrane M, Martin MJ, Natale DA, O’Donovan C, Redaschi N, Yeh L-SL. 2004. UniProt: the Universal Protein knowledgebase. Nucleic Acids Res 32:D115–9. doi:10.1093/nar/gkh131

Barlow KA, Ó Conchúir S, Thompson S, Suresh P, Lucas JE, Heinonen M, Kortemme T. 2018. Flex ddG: Rosetta Ensemble-Based Estimation of Changes in Protein-Protein Binding Affinity upon Mutation. J Phys Chem B. doi:10.1021/acs.jpcb.7b11367

Barrera LA, Vedenko A, Kurland J V, Rogers JM, Gisselbrecht SS, Rossin EJ, Woodard J, Mariani L, Kock KH, Inukai S, Siggers T, Shokri L, Gordân R, Sahni N, Cotsapas C, Hao T, Yi S, Kellis M, Daly MJ, Vidal M, Hill DE, Bulyk ML. 2016. Survey of variation in human transcription factors reveals prevalent DNA binding changes. Science 351:1450–1454. doi:10.1126/science.aad2257

Ben-Nissan G, Chotiner A, Tarnavsky M, Sharon M. 2016. Structural Characterization of Missense Mutations Using High Resolution Mass Spectrometry: A Case Study of the Parkinson’s-Related Protein, DJ-1. J Am Soc Mass Spectrom 27:1062–70. doi:10.1007/s13361-016-1379-z

Berman HM, Coimbatore Narayanan B, Di Costanzo L, Dutta S, Ghosh S, Hudson BP, Lawson CL, Peisach E, Prlić A, Rose PW, Shao C, Yang H, Young J, Zardecki C. 2013. Trendspotting in the Protein Data Bank. FEBS Lett 587:1036–45. doi:10.1016/j.febslet.2012.12.029

Berman HM, Westbrook J, Feng Z, Gilliland G, Bhat TN, Weissig H, Shindyalov IN, Bourne PE. 2000. The Protein Data Bank. Nucleic Acids Res 28:235–42.

Betz SF. 1993. Disulfide bonds and the stability of globular proteins. Protein Sci 2:1551–8. doi:10.1002/pro.5560021002

Bhattacharya R, Rose PW, Burley SK, Prlić A. 2017. Impact of genetic variation on three dimensional structure and function of proteins. PLoS One 12. doi:10.1371/journal.pone.0171355

Blundell TL. 2017. Protein crystallography and drug discovery: recollections of knowledge exchange between academia and industry. IUCrJ 4:308–321. doi:10.1107/S2052252517009241

Bonsor DA, Günther S, Beadenkopf R, Beckett D, Sundberg EJ. 2015. Diverse oligomeric states of CEACAM IgV domains. Proc Natl Acad Sci 112:13561–6.

Browning SR, Weir BS. 2010. Population structure with localized haplotype clusters. Genetics 185:1337–44. doi:10.1534/genetics.110.116681

Bycroft C, Freeman C, Petkova D, Band G, Elliott LT, Sharp K, Motyer A, Vukcevic D, Delaneau O, O’Connell J, Cortes A, Welsh S, McVean G, Leslie S, Donnelly P, Marchini J. 2017. Genome-wide genetic data on ~500,000 UK Biobank participants. bioRxiv 166298. doi:10.1101/166298

Canela-Xandri O, Rawlik K, Tenesa A. 2018. An atlas of genetic associations in UK Biobank. Nat Genet 50:1593–1599. doi:10.1038/s41588-018-0248-z

Carlson CB, Liu Y, Keck JL, Mosher DF. 2008. Influences of the N700S thrombospondin-1 polymorphism on protein structure and stability. J Biol Chem 283:20069–76. doi:10.1074/jbc.M800223200

Carlson CS, Matise TC, North KE, Haiman CA, Fesinmeyer MD, Buyske S, Schumacher FR, Peters U, Franceschini N, Ritchie MD, Duggan DJ, Spencer KL, Dumitrescu L, Eaton CB, Thomas F, Young A, Carty C, Heiss G, Le Marchand L, Crawford DC, Hindorff LA, Kooperberg CL, PAGE Consortium. 2013. Generalization and dilution of association results from European GWAS in populations of non-European ancestry: the PAGE study. PLoS Biol 11:e1001661. doi:10.1371/journal.pbio.1001661

Chan SL, Samaranayake N, Ross CJD, Toh MT, Carleton B, Hayden MR, Teo YY, Dissanayake VHW, Brunham LR. 2016. Genetic diversity of variants involved in drug response and metabolism in Sri Lankan populations: implications for clinical implementation of pharmacogenomics. Pharmacogenet Genomics 26:28–39. doi:10.1097/FPC.0000000000000182

Choi Y, Chan AP. 2015. PROVEAN web server: a tool to predict the functional effect of amino acid substitutions and indels. Bioinformatics 31:2745–7. doi:10.1093/bioinformatics/btv195

Chothia C, Lesk AM. 1986. The relation between the divergence of sequence and structure in proteins. EMBO J 5:823–6. doi:060 fehlt

Chun S, Fay JC. 2009. Identification of deleterious mutations within three human genomes. Genome Res 19:1553–61. doi:10.1101/gr.092619.109

Cline MS, Karchin R. 2010. Using bioinformatics to predict the functional impact of SNVs. Bioinformatics 27:441–448.

Cock PJA, Antao T, Chang JT, Chapman BA, Cox CJ, Dalke A, Friedberg I, Hamelryck T, Kauff F, Wilczynski B, de Hoon MJL. 2009. Biopython: freely available Python tools for computational molecular biology and bioinformatics. Bioinformatics 25:1422–1423. doi:10.1093/bioinformatics/btp163

Cohen P. 2001. The role of protein phosphorylation in human health and disease. The Sir Hans Krebs Medal Lecture. Eur J Biochem 268:5001–10.

Consortium GLG, Willer CJ, Schmidt EM, Sengupta S, Peloso GM, Gustafsson S, Kanoni S, Ganna A, Chen J, Buchkovich ML, Mora S, Beckmann JS, Bragg-Gresham JL, Chang H-Y, Demirkan A, Den Hertog HM, Do R, Donnelly LA, Ehret GB, Esko T, Feitosa MF, Ferreira T, Fischer K, Fontanillas P, Fraser RM, Freitag DF, Gurdasani D, Heikkilä K, Hyppönen E, Isaacs A, Jackson AU, Johansson Å, Johnson T, Kaakinen M, Kettunen J, Kleber ME, Li X, Luan J, Lyytikäinen L-P, Magnusson PKE, Mangino M, Mihailov E, Montasser ME, Müller-Nurasyid M, Nolte IM, O’Connell JR, Palmer CD, Perola M, Petersen A-K, Sanna S, Saxena R, Service SK, Shah S, Shungin D, Sidore C, Song C, Strawbridge RJ, Surakka I, Tanaka T, Teslovich TM, Thorleifsson G, Van den Herik EG, Voight BF, Volcik KA, Waite LL, Wong A, Wu Y, Zhang W, Absher D, Asiki G, Barroso I, Been LF, Bolton JL, Bonnycastle LL, Brambilla P, Burnett MS, Cesana G, Dimitriou M, Doney ASF, Döring A, Elliott P, Epstein SE, Eyjolfsson GI, Gigante B, Goodarzi MO, Grallert H, Gravito ML, Groves CJ, Hallmans G, Hartikainen A-L, Hayward C, Hernandez D, Hicks AA, Holm H, Hung Y-J, Illig T, Jones MR, Kaleebu P, Kastelein JJP, Khaw K-T, Kim E, Klopp N, Komulainen P, Kumari M, Langenberg C, Lehtimäki T, Lin S-Y, Lindström J, Loos RJF, Mach F, McArdle WL, Meisinger C, Mitchell BD, Müller G, Nagaraja R, Narisu N, Nieminen TVM, Nsubuga RN, Olafsson I, Ong KK, Palotie A, Papamarkou T, Pomilla C, Pouta A, Rader DJ, Reilly MP, Ridker PM, Rivadeneira F, Rudan I, Ruokonen A, Samani N, Scharnagl H, Seeley J, Silander K, Stancáková A, Stirrups K, Swift AJ, Tiret L, Uitterlinden AG, van Pelt LJ, Vedantam S, Wainwright N, Wijmenga C, Wild SH, Willemsen G, Wilsgaard T, Wilson JF, Young EH, Zhao JH, Adair LS, Arveiler D, Assimes TL, Bandinelli S, Bennett F, Bochud M, Boehm BO, Boomsma DI, Borecki IB, Bornstein SR, Bovet P, Burnier M, Campbell H, Chakravarti A, Chambers JC, Chen Y-DI, Collins FS, Cooper RS, Danesh J, Dedoussis G, de Faire U, Feranil AB, Ferrières J, Ferrucci L, Freimer NB, Gieger C, Groop LC, Gudnason V, Gyllensten U, Hamsten A, Harris TB, Hingorani A, Hirschhorn JN, Hofman A, Hovingh GK, Hsiung CA, Humphries SE, Hunt SC, Hveem K, Iribarren C, Järvelin M-R, Jula A, Kähönen M, Kaprio J, Kesäniemi A, Kivimaki M, Kooner JS, Koudstaal PJ, Krauss RM, Kuh D, Kuusisto J, Kyvik KO, Laakso M, Lakka TA, Lind L, Lindgren CM, Martin NG, März W, McCarthy MI, McKenzie CA, Meneton P, Metspalu A, Moilanen L, Morris AD, Munroe PB, Njølstad I, Pedersen NL, Power C, Pramstaller PP, Price JF, Psaty BM, Quertermous T, Rauramaa R, Saleheen D, Salomaa V, Sanghera DK, Saramies J, Schwarz PEH, Sheu WH-H, Shuldiner AR, Siegbahn A, Spector TD, Stefansson K, Strachan DP, Tayo BO, Tremoli E, Tuomilehto J, Uusitupa M, van Duijn CM, Vollenweider P, Wallentin L, Wareham NJ, Whitfield JB, Wolffenbuttel BHR, Ordovas JM, Boerwinkle E, Palmer CNA, Thorsteinsdottir U, Chasman DI, Rotter JI, Franks PW, Ripatti S, Cupples LA, Sandhu MS, Rich SS, Boehnke M, Deloukas P, Kathiresan S, Mohlke KL, Ingelsson E, Abecasis GR. 2013. Discovery and refinement of loci associated with lipid levels. Nat Genet 45:1274.

Cooper DN, Chen J-M, Ball E V, Howells K, Mort M, Phillips AD, Chuzhanova N, Krawczak M, Kehrer-Sawatzki H, Stenson PD. 2010. Genes, mutations, and human inherited disease at the dawn of the age of personalized genomics. Hum Mutat 31:631–55. doi:10.1002/humu.21260

Crawford NG, Kelly DE, Hansen MEB, Beltrame MH, Fan S, Bowman SL, Jewett E, Ranciaro A, Thompson S, Lo Y, Pfeifer SP, Jensen JD, Campbell MC, Beggs W, Hormozdiari F, Mpoloka SW, Mokone GG, Nyambo T, Meskel DW, Belay G, Haut J, Rothschild H, Zon L, Zhou Y, Kovacs MA, Xu M, Zhang T, Bishop K, Sinclair J, Rivas C, Elliot E, Choi J, Li SA, Hicks B, Burgess S, Abnet C, Watkins-Chow DE, Oceana E, Song YS, Eskin E, Brown KM, Marks MS, Loftus SK, Pavan WJ, Yeager M, Chanock S, Tishkoff SA. 2017. Loci associated with skin pigmentation identified in African populations. Science 358. doi:10.1126/science.aan8433

Creixell P, Schoof EM, Simpson CD, Longden J, Miller CJ, Lou HJ, Perryman L, Cox TR, Zivanovic N, Palmeri A, Wesolowska-Andersen A, Helmer-Citterich M, Ferkinghoff-Borg J, Itamochi H, Bodenmiller B, Erler JT, Turk BE, Linding R. 2015. Kinome-wide decoding of network-attacking mutations rewiring cancer signaling. Cell 163:202–17. doi:10.1016/j.cell.2015.08.056

Davis AM, St-Gallay SA, Kleywegt GJ. 2008. Limitations and lessons in the use of X-ray structural information in drug design. Drug Discov Today 13:831–41. doi:10.1016/j.drudis.2008.06.006

de Beer TAP, Laskowski RA, Parks SL, Sipos B, Goldman N, Thornton JM. 2013. Amino acid changes in disease-associated variants differ radically from variants observed in the 1000 genomes project dataset. PLoS Comput Biol 9:e1003382.

Dey S, Ritchie DW, Levy ED. 2018. PDB-wide identification of biological assemblies from conserved quaternary structure geometry. Nat Methods 15:67–72. doi:10.1038/nmeth.4510

Dong C, Wei P, Jian X, Gibbs R, Boerwinkle E, Wang K, Liu X. 2015. Comparison and integration of deleteriousness prediction methods for nonsynonymous SNVs in whole exome sequencing studies. Hum Mol Genet 24:2125–37. doi:10.1093/hmg/ddu733

Ely ZA, Moon JM, Sliwoski GR, Sangha AK, Shen X-X, LaBella AL, Meiler J, Capra JA, Rokas A. 2018. The impact of natural selection on the evolution and function of placentally expressed galectins. bioRxiv 505339. doi:10.1101/505339

Fan S, Hansen MEB, Lo Y, Tishkoff SA. 2016. Going global by adapting local: A review of recent human adaptation. Science 354:54–59. doi:10.1126/science.aaf5098

Ferrer-Costa C, Orozco M, de la Cruz X. 2002. Characterization of disease-associated single amino acid polymorphisms in terms of sequence and structure properties. J Mol Biol 315:771–86. doi:10.1006/jmbi.2001.5255

Fumagalli M, Moltke I, Grarup N, Racimo F, Bjerregaard P, Jørgensen ME, Korneliussen TS, Gerbault P, Skotte L, Linneberg A, Christensen C, Brandslund I, Jørgensen T, Huerta-Sánchez E, Schmidt EB, Pedersen O, Hansen T, Albrechtsen A, Nielsen R. 2015. Greenlandic Inuit show genetic signatures of diet and climate adaptation. Science 349:1343–7. doi:10.1126/science.aab2319

Gao M, Zhou H, Skolnick J. 2015. Insights into disease-associated mutations in the human proteome through protein structural analysis. Structure 23:1362–1369.

Hauser AS, Chavali S, Masuho I, Jahn LJ, Martemyanov KA, Gloriam DE, Babu MM. 2018. Pharmacogenomics of GPCR Drug Targets. Cell 172:41-54.e19. doi:10.1016/j.cell.2017.11.033

Hindorff LA, Bonham VL, Brody LC, Ginoza MEC, Hutter CM, Manolio TA, Green ED. 2018. Prioritizing diversity in human genomics research. Nat Rev Genet 19:175–185. doi:10.1038/nrg.2017.89

Humphrey SJ, James DE, Mann M. 2015. Protein Phosphorylation: A Major Switch Mechanism for Metabolic Regulation. Trends Endocrinol Metab 26:676–687. doi:10.1016/j.tem.2015.09.013

Iotchkova V, Huang J, Morris JA, Jain D, Barbieri C, Walter K, Min JL, Chen L, Astle W, Cocca M, Deelen P, Elding H, Farmaki A-E, Franklin CS, Franberg M, Gaunt TR, Hofman A, Jiang T, Kleber ME, Lachance G, Luan J, Malerba G, Matchan A, Mead D, Memari Y, Ntalla I, Panoutsopoulou K, Pazoki R, Perry JRB, Rivadeneira F, Sabater-Lleal M, Sennblad B, Shin S-Y, Southam L, Traglia M, van Dijk F, van Leeuwen EM, Zaza G, Zhang W, Consortium U, Amin N, Butterworth A, Chambers JC, Dedoussis G, Dehghan A, Franco OH, Franke L, Frontini M, Gambaro G, Gasparini P, Hamsten A, Issacs A, Kooner JS, Kooperberg C, Langenberg C, Marz W, Scott RA, Swertz MA, Toniolo D, Uitterlinden AG, van Duijn CM, Watkins H, Zeggini E, Maurano MT, Timpson NJ, Reiner AP, Auer PL, Soranzo N. 2016. Discovery and refinement of genetic loci associated with cardiometabolic risk using dense imputation maps. Nat Genet 48:1303.

Ishchenko A, Gati C, Cherezov V. 2018. Structural biology of G protein-coupled receptors: new opportunities from XFELs and cryoEM. Curr Opin Struct Biol 51:44–52. doi:10.1016/j.sbi.2018.03.009

Jeong DG, Cho YH, Yoon T-S, Kim JH, Ryu SE, Kim SJ. 2007. Crystal structure of the catalytic domain of human DUSP5, a dual specificity MAP kinase protein phosphatase. Proteins 66:253–8. doi:10.1002/prot.21224

Johansson KE, Lindorff-Larsen K. 2018. Structural heterogeneity and dynamics in protein evolution and design. Curr Opin Struct Biol 48:157–163. doi:10.1016/j.sbi.2018.01.010

Kang HJ, Wilkins AD, Lichtarge O, Wensel TG. 2015. Determinants of endogenous ligand specificity divergence among metabotropic glutamate receptors. J Biol Chem 290:2870–8. doi:10.1074/jbc.M114.622233

Kann MG. 2007. Protein interactions and disease: computational approaches to uncover the etiology of diseases. Brief Bioinform 8:333–46. doi:10.1093/bib/bbm031

Kellogg EH, Leaver-Fay A, Baker D. 2011. Role of conformational sampling in computing mutation-induced changes in protein structure and stability. Proteins 79:830–8. doi:10.1002/prot.22921

Kessler MD, Yerges-Armstrong L, Taub MA, Shetty AC, Maloney K, Jeng LJB, Ruczinski I, Levin AM, Williams LK, Beaty TH, Mathias RA, Barnes KC, Boorgula MP, Campbell M, Chavan S, Ford JG, Foster C, Gao L, Hansel NN, Horowitz E, Huang L, Ortiz R, Potee J, Rafaels N, Scott AF, Vergara C, Gao J, Hu Y, Johnston HR, Qin ZS, Padhukasahasram B, Dunston GM, Faruque MU, Kenny EE, Gietzen K, Hansen M, Genuario R, Bullis D, Lawley C, Deshpande A, Grus WE, Locke DP, Foreman MG, Avila PC, Grammer L, Kim K-Y, Kumar R, Schleimer R, Bustamante C, De La Vega FM, Gignoux CR, Shringarpure SS, Musharoff S, Wojcik G, Burchard EG, Eng C, Gourraud P-A, Hernandez RD, Lizee A, Pino-Yanes M, Torgerson DG, Szpiech ZA, Torres R, Nicolae DL, Ober C, Olopade CO, Olopade O, Oluwole O, Arinola G, Song W, Abecasis G, Correa A, Musani S, Wilson JG, Lange LA, Akey J, Bamshad M, Chong J, Fu W, Nickerson D, Reiner A, Hartert T, Ware LB, Bleecker E, Meyers D, Ortega VE, Pissamai MRN, Trevor MRN, Watson H, Araujo MI, Oliveira RR, Caraballo L, Marrugo J, Martinez B, Meza C, Ayestas G, Herrera-Paz EF, Landaverde-Torres P, Erazo SOL, Martinez R, Mayorga A, Mayorga LF, Mejia-Mejia D-A, Ramos H, Saenz A, Varela G, Vasquez OM, Ferguson T, Knight-Madden J, Samms-Vaughan M, Wilks RJ, Adegnika A, Ateba-Ngoa U, Yazdanbakhsh M, O’Connor TD, O’Connor TD. 2016. Challenges and disparities in the application of personalized genomic medicine to populations with African ancestry. Nat Commun 7:12521. doi:10.1038/ncomms12521

Khoo KK, Norton RS. 2011. Role of Disulfide Bonds in Peptide and Protein ConformationAmino Acids, Peptides and Proteins in Organic Chemistry. Weinheim, Germany: Wiley-VCH Verlag GmbH & Co. KGaA. pp. 395–417. doi:10.1002/9783527631841.ch11

Kumari V, Dyba MA, Holland RJ, Liang Y-H, Singh S V, Ji X. 2016. Irreversible Inhibition of Glutathione S-Transferase by Phenethyl Isothiocyanate (PEITC), a Dietary Cancer Chemopreventive Phytochemical. PLoS One 11:1–12. doi:10.1371/journal.pone.0163821

Law V, Knox C, Djoumbou Y, Jewison T, Guo AC, Liu Y, Maciejewski A, Arndt D, Wilson M, Neveu V, Tang A, Gabriel G, Ly C, Adamjee S, Dame ZT, Han B, Zhou Y, Wishart DS. 2014. DrugBank 4.0: shedding new light on drug metabolism. Nucleic Acids Res 42:D1091–7. doi:10.1093/nar/gkt1068

Lek M, Karczewski KJ, Minikel E V, Samocha KE, Banks E, Fennell T, O’Donnell-Luria AH, Ware JS, Hill AJ, Cummings BB, Tukiainen T, Birnbaum DP, Kosmicki JA, Duncan LE, Estrada K, Zhao F, Zou J, Pierce-Hoffman E, Berghout J, Cooper DN, Deflaux N, DePristo M, Do R, Flannick J, Fromer M, Gauthier L, Goldstein J, Gupta N, Howrigan D, Kiezun A, Kurki MI, Moonshine AL, Natarajan P, Orozco L, Peloso GM, Poplin R, Rivas MA, Ruano-Rubio V, Rose SA, Ruderfer DM, Shakir K, Stenson PD, Stevens C, Thomas BP, Tiao G, Tusie-Luna MT, Weisburd B, Won H-H, Yu D, Altshuler DM, Ardissino D, Boehnke M, Danesh J, Donnelly S, Elosua R, Florez JC, Gabriel SB, Getz G, Glatt SJ, Hultman CM, Kathiresan S, Laakso M, McCarroll S, McCarthy MI, McGovern D, McPherson R, Neale BM, Palotie A, Purcell SM, Saleheen D, Scharf JM, Sklar P, Sullivan PF, Tuomilehto J, Tsuang MT, Watkins HC, Wilson JG, Daly MJ, MacArthur DG, Exome Aggregation Consortium. 2016. Analysis of protein-coding genetic variation in 60,706 humans. Nature 536:285–91. doi:10.1038/nature19057

Levy ED. 2007. PiQSi: protein quaternary structure investigation. Structure 15:1364–7. doi:10.1016/j.str.2007.09.019

Li Y, Yan J, Zhang X, Huang K. 2013. Disulfide bonds in amyloidogenesis diseases related proteins. Proteins 81:1862–73. doi:10.1002/prot.24338

Li YI, van de Geijn B, Raj A, Knowles DA, Petti AA, Golan D, Gilad Y, Pritchard JK. 2016. RNA splicing is a primary link between genetic variation and disease. Science 352:600–4. doi:10.1126/science.aad9417

Lindner I, Rubin D, Helwig U, Nitz I, Hampe J, Schreiber S, Schrezenmeir J, Döring F. 2006. The L513S polymorphism in medium-chain acyl-CoA synthetase 2 (MACS2) is associated with risk factors of the metabolic syndrome in a Caucasian study population. Mol Nutr Food Res 50:270–4. doi:10.1002/mnfr.200500241

Liu X, Jian X, Boerwinkle E. 2011. dbNSFP: a lightweight database of human nonsynonymous SNPs and their functional predictions. Hum Mutat 32:894–9. doi:10.1002/humu.21517

Liu Z, Li Y, Han L, Li J, Liu J, Zhao Z, Nie W, Liu Y, Wang R. 2015. PDB-wide collection of binding data: current status of the PDBbind database. Bioinformatics 31:405–12. doi:10.1093/bioinformatics/btu626

Lu H-C, Fornili A, Fraternali F. 2013. Protein-protein interaction networks studies and importance of 3D structure knowledge. Expert Rev Proteomics 10:511–20. doi:10.1586/14789450.2013.856764

Magliery TJ, Regan L. 2005. Sequence variation in ligand binding sites in proteins. BMC Bioinformatics 6:240. doi:10.1186/1471-2105-6-240

Mahmood K, Jung C, Philip G, Georgeson P, Chung J, Pope BJ, Park DJ. 2017. Variant effect prediction tools assessed using independent, functional assay-based datasets: implications for discovery and diagnostics. Hum Genomics 11:10. doi:10.1186/s40246-017-0104-8

Mallick S, Li H, Lipson M, Mathieson I, Gymrek M, Racimo F, Zhao M, Chennagiri N, Nordenfelt S, Tandon A, Skoglund P, Lazaridis I, Sankararaman S, Fu Q, Rohland N, Renaud G, Erlich Y, Willems T, Gallo C, Spence JP, Song YS, Poletti G, Balloux F, Van Driem G, De Knijff P, Romero IG, Jha AR, Behar DM, Bravi CM, Capelli C, Hervig T, Moreno-Estrada A, Posukh OL, Balanovska E, Balanovsky O, Karachanak-Yankova S, Sahakyan H, Toncheva D, Yepiskoposyan L, Tyler-Smith C, Xue Y, Abdullah MS, Ruiz-Linares A, Beall CM, Di Rienzo A, Jeong C, Starikovskaya EB, Metspalu E, Parik J, Villems R, Henn BM, Hodoglugil U, Mahley R, Sajantila A, Stamatoyannopoulos G, Wee JTS, Khusainova R, Khusnutdinova E, Litvinov S, Ayodo G, Comas D, Hammer MF, Kivisild T, Klitz W, Winkler CA, Labuda D, Bamshad M, Jorde LB, Tishkoff SA, Watkins WS, Metspalu M, Dryomov S, Sukernik R, Singh L, Thangaraj K, Paäbo S, Kelso J, Patterson N, Reich D. 2016. The Simons Genome Diversity Project: 300 genomes from 142 diverse populations. Nature 538:201–206. doi:10.1038/nature18964

Mannige R V. 2014. Dynamic New World: Refining Our View of Protein Structure, Function and Evolution. Proteomes 2:128–153. doi:10.3390/proteomes2010128

Manrai AK, Funke BH, Rehm HL, Olesen MS, Maron BA, Szolovits P, Margulies DM, Loscalzo J, Kohane IS. 2016. Genetic Misdiagnoses and the Potential for Health Disparities. N Engl J Med 375:655–65. doi:10.1056/NEJMsa1507092

Martelli PL, Fariselli P, Savojardo C, Babbi G, Aggazio F, Casadio R. 2016. Large scale analysis of protein stability in OMIM disease related human protein variants. BMC Genomics 17 Suppl 2:397. doi:10.1186/s12864-016-2726-y

Mathias RA, Taub MA, Gignoux CR, Fu W, Musharoff S, O’Connor TD, Vergara C, Torgerson DG, Pino-Yanes M, Shringarpure SS, Huang L, Rafaels N, Boorgula MP, Johnston HR, Ortega VE, Levin AM, Song W, Torres R, Padhukasahasram B, Eng C, Mejia-Mejia D-A, Ferguson T, Qin ZS, Scott AF, Yazdanbakhsh M, Wilson JG, Marrugo J, Lange LA, Kumar R, Avila PC, Williams LK, Watson H, Ware LB, Olopade C, Olopade O, Oliveira R, Ober C, Nicolae DL, Meyers D, Mayorga A, Knight-Madden J, Hartert T, Hansel NN, Foreman MG, Ford JG, Faruque MU, Dunston GM, Caraballo L, Burchard EG, Bleecker E, Araujo MI, Herrera-Paz EF, Gietzen K, Grus WE, Bamshad M, Bustamante CD, Kenny EE, Hernandez RD, Beaty TH, Ruczinski I, Akey J, Caapa, Barnes KC. 2016. A continuum of admixture in the Western Hemisphere revealed by the African Diaspora genome. Nat Commun 7:12522. doi:10.1038/ncomms12522

McCarthy DJ, Humburg P, Kanapin A, Rivas MA, Gaulton K, Cazier J-B, Donnelly P. 2014. Choice of transcripts and software has a large effect on variant annotation. Genome Med 6:26. doi:10.1186/gm543

McLaren W, Gil L, Hunt SE, Riat HS, Ritchie GRS, Thormann A, Flicek P, Cunningham F. 2016. The Ensembl Variant Effect Predictor. Genome Biol 17:122. doi:10.1186/s13059-016-0974-4

Mungall C, fbastian, Lewis S, smanzoor, kltm, dougli1sqrd, hdietze, tudorgroza, Overton JA, Keith D, Himmelstein D. 2017. owlcollab/owltools: v0.3.0. doi:10.5281/ZENODO.574119

Nei M, Tajima F. 1981. DNA polymorphism detectable by restriction endonucleases. Genetics 97:145–63.

Nelson MR, Wegmann D, Ehm MG, Kessner D, St Jean P, Verzilli C, Shen J, Tang Z, Bacanu S-A, Fraser D, Warren L, Aponte J, Zawistowski M, Liu X, Zhang H, Zhang Y, Li J, Li Y, Li L, Woollard P, Topp S, Hall MD, Nangle K, Wang J, Abecasis G, Cardon LR, Zöllner S, Whittaker JC, Chissoe SL, Novembre J, Mooser V. 2012. An abundance of rare functional variants in 202 drug target genes sequenced in 14,002 people. Science 337:100–4. doi:10.1126/science.1217876

Ng PC, Henikoff S. 2001. Predicting deleterious amino acid substitutions. Genome Res 11:863–74. doi:10.1101/gr.176601

Nielsen R, Akey JM, Jakobsson M, Pritchard JK, Tishkoff S, Willerslev E. 2017. Tracing the peopling of the world through genomics. Nature 541:302–310. doi:10.1038/nature21347

Nishio K, Yoshida Y, Tanaka K, Mizushima T. 2016. Structural analysis of a function-associated loop mutant of the substrate-recognition domain of Fbs1 ubiquitin ligase. Acta Crystallogr Sect F, Struct Biol Commun 72:619–26. doi:10.1107/S2053230X16011018

Ongen H, Dermitzakis ET. 2015. Alternative Splicing QTLs in European and African Populations. Am J Hum Genet 97:567–75. doi:10.1016/j.ajhg.2015.09.004

Park E, Pan Z, Zhang Z, Lin L, Xing Y. 2018. The Expanding Landscape of Alternative Splicing Variation in Human Populations. Am J Hum Genet 102:11–26. doi:10.1016/j.ajhg.2017.11.002

Perera MA, Cavallari LH, Johnson JA. 2014. Warfarin pharmacogenetics: an illustration of the importance of studies in minority populations. Clin Pharmacol Ther 95:242–4. doi:10.1038/clpt.2013.209

Petrovski S, Goldstein DB. 2016. Unequal representation of genetic variation across ancestry groups creates healthcare inequality in the application of precision medicine. Genome Biol 17:157. doi:10.1186/s13059-016-1016-y

Pettersen EF, Goddard TD, Huang CC, Couch GS, Greenblatt DM, Meng EC, Ferrin TE. 2004. UCSF Chimera--a visualization system for exploratory research and analysis. J Comput Chem 25:1605–12. doi:10.1002/jcc.20084

Petukh M, Kucukkal TG, Alexov E. 2015. On human disease-causing amino acid variants: statistical study of sequence and structural patterns. Hum Mutat 36:524–534. doi:10.1002/humu.22770

Popejoy AB, Fullerton SM. 2016. Genomics is failing on diversity. Nature 538:161–164. doi:10.1038/538161a

Preissner SC, Hoffmann MF, Preissner R, Dunkel M, Gewiess A, Preissner S. 2013. Polymorphic cytochrome P450 enzymes (CYPs) and their role in personalized therapy. PLoS One 8:e82562. doi:10.1371/journal.pone.0082562

Prlic A, Kalro T, Bhattacharya R, Christie C, Burley SK, Rose PW. 2016. Integrating genomic information with protein sequence and 3D atomic level structure at the RCSB protein data bank. Bioinformatics. doi:10.1093/bioinformatics/btw547

Roden DM. 2016. Cardiovascular pharmacogenomics: current status and future directions. J Hum Genet 61:79–85. doi:10.1038/jhg.2015.78

Ruderfer DM, Hamamsy T, Lek M, Karczewski KJ, Kavanagh D, Samocha KE, Exome Aggregation Consortium, Daly MJ, MacArthur DG, Fromer M, Purcell SM. 2016. Patterns of genic intolerance of rare copy number variation in 59,898 human exomes. Nat Genet 48:1107–11. doi:10.1038/ng.3638

Ryan M, Diekhans M, Lien S, Liu Y, Karchin R. 2009. LS-SNP/PDB: annotated non-synonymous SNPs mapped to Protein Data Bank structures. Bioinformatics 25:1431–2. doi:10.1093/bioinformatics/btp242

Sanghera DK, Wagenknecht DR, McIntyre JA, Kamboh MI. 1997. Identification of structural mutations in the fifth domain of apolipoprotein H (beta 2-glycoprotein I) which affect phospholipid binding. Hum Mol Genet 6:311–6.

Sauna ZE, Kimchi-Sarfaty C. 2011. Understanding the contribution of synonymous mutations to human disease. Nat Rev Genet 12:683–91. doi:10.1038/nrg3051

Scott DE, Bayly AR, Abell C, Skidmore J. 2016. Small molecules, big targets: drug discovery faces the protein-protein interaction challenge. Nat Rev Drug Discov 15:533–50. doi:10.1038/nrd.2016.29

Seeger MA. 2018. Membrane transporter research in times of countless structures. Biochim Biophys Acta 1860:804–808. doi:10.1016/j.bbamem.2017.08.009

Shihab HA, Gough J, Cooper DN, Stenson PD, Barker GLA, Edwards KJ, Day INM, Gaunt TR. 2013. Predicting the functional, molecular, and phenotypic consequences of amino acid substitutions using hidden Markov models. Hum Mutat 34:57–65. doi:10.1002/humu.22225

Sivley RM, Dou X, Meiler J, Bush WS, Capra JA. 2018. Comprehensive Analysis of Constraint on the Spatial Distribution of Missense Variants in Human Protein Structures. Am J Hum Genet 102:415–426. doi:10.1016/j.ajhg.2018.01.017

Śledź P, Caflisch A. 2018. Protein structure-based drug design: from docking to molecular dynamics. Curr Opin Struct Biol 48:93–102. doi:10.1016/j.sbi.2017.10.010

Smith CA, Kortemme T. 2008. Backrub-like backbone simulation recapitulates natural protein conformational variability and improves mutant side-chain prediction. J Mol Biol 380:742–56. doi:10.1016/j.jmb.2008.05.023

Somody JC, MacKinnon SS, Windemuth A. 2017. Structural coverage of the proteome for pharmaceutical applications. Drug Discov Today 22:1792–1799. doi:10.1016/j.drudis.2017.08.004

Spooner W, McLaren W, Slidel T, Finch DK, Butler R, Campbell J, Eghobamien L, Rider D, Kiefer CM, Robinson MJ, Hardman C, Cunningham F, Vaughan T, Flicek P, Huntington CC. 2018. Haplosaurus computes protein haplotypes for use in precision drug design. Nat Commun 9:4128. doi:10.1038/s41467-018-06542-1

Suhre K, Arnold M, Bhagwat AM, Cotton RJ, Engelke R, Raffler J, Sarwath H, Thareja G, Wahl A, DeLisle RK, Gold L, Pezer M, Lauc G, El-Din Selim MA, Mook-Kanamori DO, Al-Dous EK, Mohamoud YA, Malek J, Strauch K, Grallert H, Peters A, Kastenmüller G, Gieger C, Graumann J. 2017. Connecting genetic risk to disease end points through the human blood plasma proteome. Nat Commun 8:14357.

Tang H, Thomas PD. 2016. Tools for Predicting the Functional Impact of Nonsynonymous Genetic Variation. Genetics 203:635–47. doi:10.1534/genetics.116.190033

Teilum K, Olsen JG, Kragelund BB. 2011. Protein stability, flexibility and function. Biochim Biophys Acta 1814:969–76. doi:10.1016/j.bbapap.2010.11.005

Tekola-Ayele F, Adeyemo A, Aseffa A, Hailu E, Finan C, Davey G, Rotimi CN, Newport MJ. 2015. Clinical and pharmacogenomic implications of genetic variation in a Southern Ethiopian population. Pharmacogenomics J 15:101–108. doi:10.1038/tpj.2014.39

Tekola-Ayele F, Peprah E. 2017. Examining How Our Shared Evolutionary History Shapes Future Disease Outcomes. Glob Heart 12:169–171. doi:10.1016/j.gheart.2017.01.008

Tennessen JA, Bigham AW, O’Connor TD, Fu W, Kenny EE, Gravel S, McGee S, Do R, Liu X, Jun G, Kang HM, Jordan D, Leal SM, Gabriel S, Rieder MJ, Abecasis G, Altshuler D, Nickerson DA, Boerwinkle E, Sunyaev S, Bustamante CD, Bamshad MJ, Akey JM, Broad GO, Seattle GO, NHLBI Exome Sequencing Project. 2012. Evolution and functional impact of rare coding variation from deep sequencing of human exomes. Science 337:64–9. doi:10.1126/science.1219240

Velankar S, Dana JM, Jacobsen J, van Ginkel G, Gane PJ, Luo J, Oldfield TJ, O’Donovan C, Martin M-J, Kleywegt GJ. 2013. SIFTS: Structure Integration with Function, Taxonomy and Sequences resource. Nucleic Acids Res 41:D483–9. doi:10.1093/nar/gks1258

Wang K, Li M, Hakonarson H. 2010. ANNOVAR: functional annotation of genetic variants from high-throughput sequencing data. Nucleic Acids Res 38:e164. doi:10.1093/nar/gkq603

Wang Z, Moult J. 2001. SNPs, protein structure, and disease. Hum Mutat 17:263–270.

Weir BS, Cockerham CC. 1984. Estimating F-statistics for the Analysis of Population Structure. Evolution (N Y) 38:1358–1370. doi:10.1111/j.1558-5646.1984.tb05657.x

Wishart DS, Knox C, Guo AC, Cheng D, Shrivastava S, Tzur D, Gautam B, Hassanali M. 2008. DrugBank: a knowledgebase for drugs, drug actions and drug targets. Nucleic Acids Res 36:D901–6. doi:10.1093/nar/gkm958

Zanger UM, Schwab M. 2013. Cytochrome P450 enzymes in drug metabolism: regulation of gene expression, enzyme activities, and impact of genetic variation. Pharmacol Ther 138:103–41. doi:10.1016/j.pharmthera.2012.12.007

Zeth K, Zachariae U. 2018. Ten Years of High Resolution Structural Research on the Voltage Dependent Anion Channel (VDAC)-Recent Developments and Future Directions. Front Physiol 9:108. doi:10.3389/fphys.2018.00108

