## Supplementary Information for "Do available protein 3D structures reflect human genetic and functional diversity?"

### *Supplementary Figures*

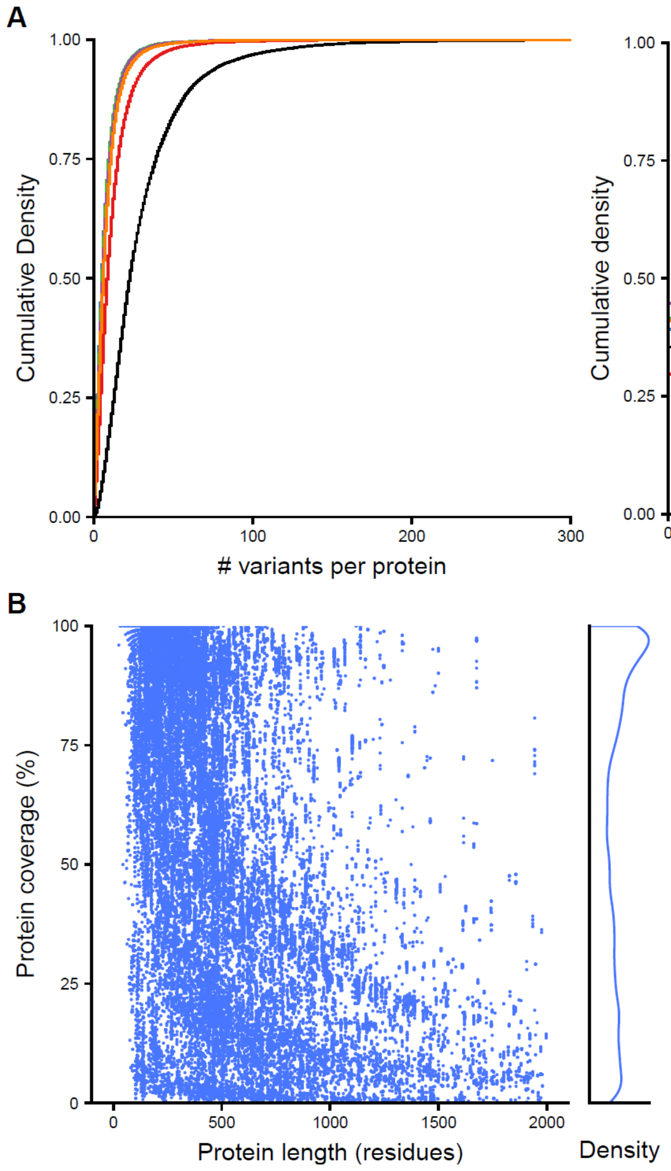


**Figure S1. Total single residue missense variants per protein in the 1000G dataset**
(A) Empirical cumulative density plot of all single residue missense variants in 1000G stratified by population. Left = all variants; right = variants with frequency >=0.01. Black = All populations; red = AFR; blue = AMR; green = EUR; purple = EAS; orange = SAS. The X-axis limits were set for visibility but 31 proteins contain more than 300 variants and 4 proteins contain more than 100 variants with frequency >=0.01. (B) The percent of each protein sequence covered by structures is plotted against the length of the protein sequence. Each point represents a single structure (chain in the PDB). The right margin plots the distribution of the protein coverage by structures.

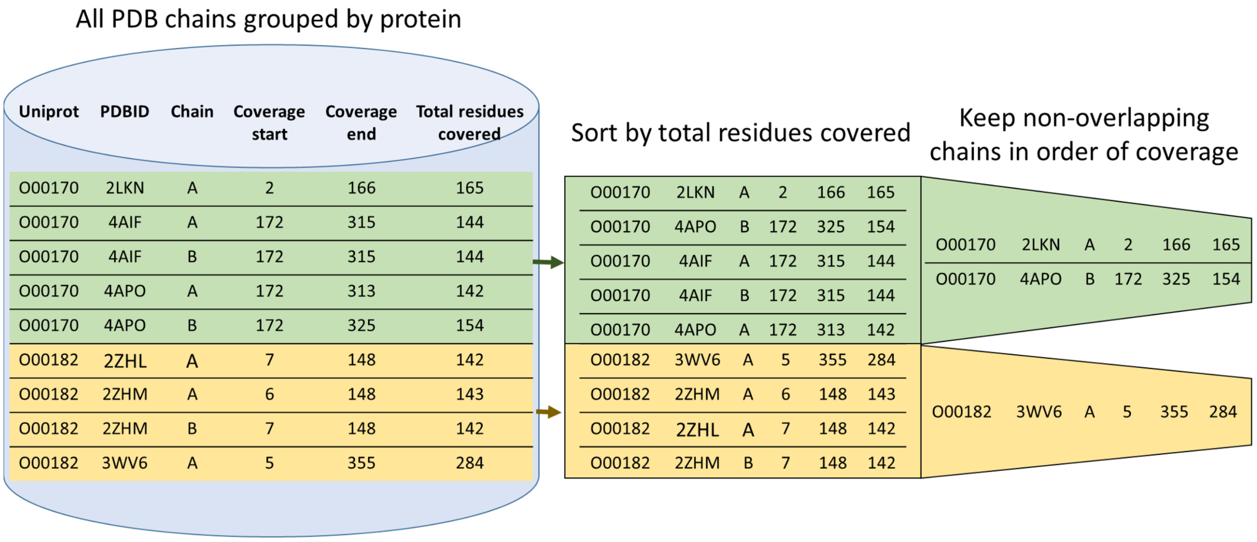


**Figure S2. Non-redundant chain selection pipeline**

The pipeline for selecting non-redundant protein chains from the PDB consists of three steps: 1. Group all PDB models by their UniProt identifier. 2. Sort each group by the total number of transcript residues covered by each PDB model. 3. Beginning with the model covering the most residues, keep each model that does not overlap with a previously kept model by 50% or more.

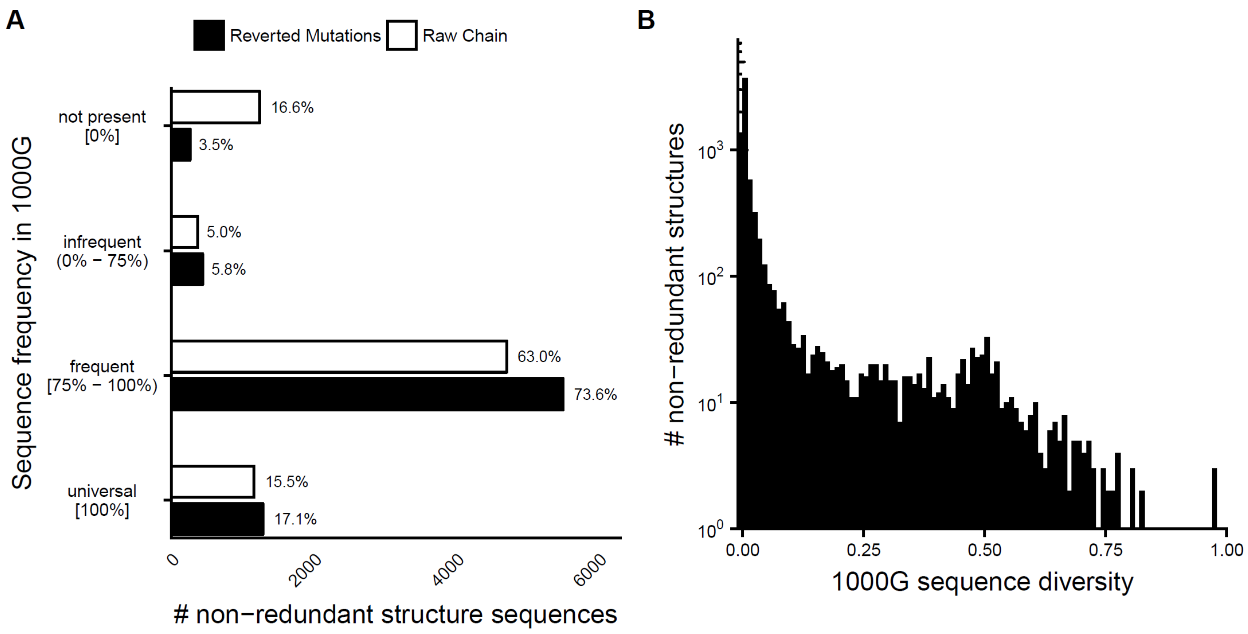


**Figure S3. Protein sequences represented in the non-redundant set of structures do not match the sequences found in many individuals.** (A) Version of Main Text Figure 2A based on the non-redundant set of structures. Histograms of the frequency across individuals in 1000G of each sequence represented for human protein structures in the non-redundant set. The white bars reflect the frequencies of the raw sequence in the protein structure. The black bars show the sequence frequencies after reverting engineered mutations in the protein structure to the original transcript amino acid. The structures are binned based on their frequency across 1000G. (B) Version of Main Text Figure 2B based on non-redundant structures. Histogram of sequence haplotype diversity computed across 1000G individuals for sequences covered by human PDB structures. Thousands of proteins represented by structures have high levels of diversity across human populations.

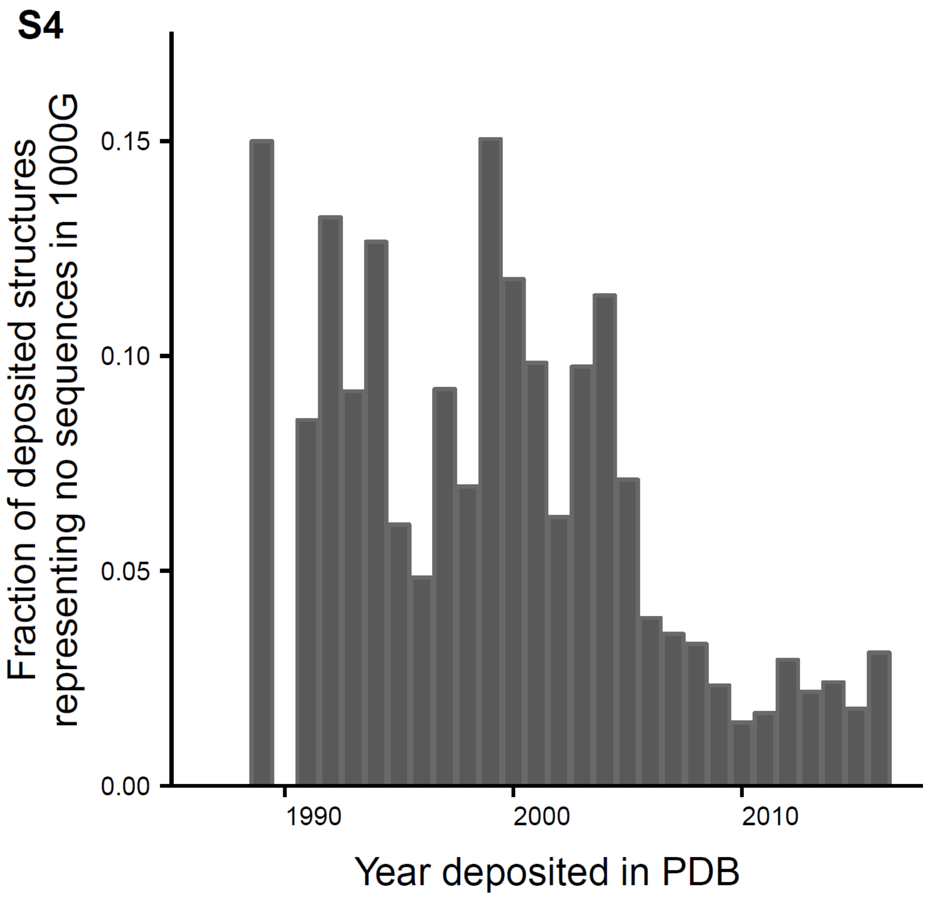


**Figure S4. Human structures deposited in the PDB since 2005 are less likely not to match a sequence observed in 1000G.** Histogram of the fraction of chains deposited in the PDB each year since 1985 that do not match any sequence observed in 1000G.

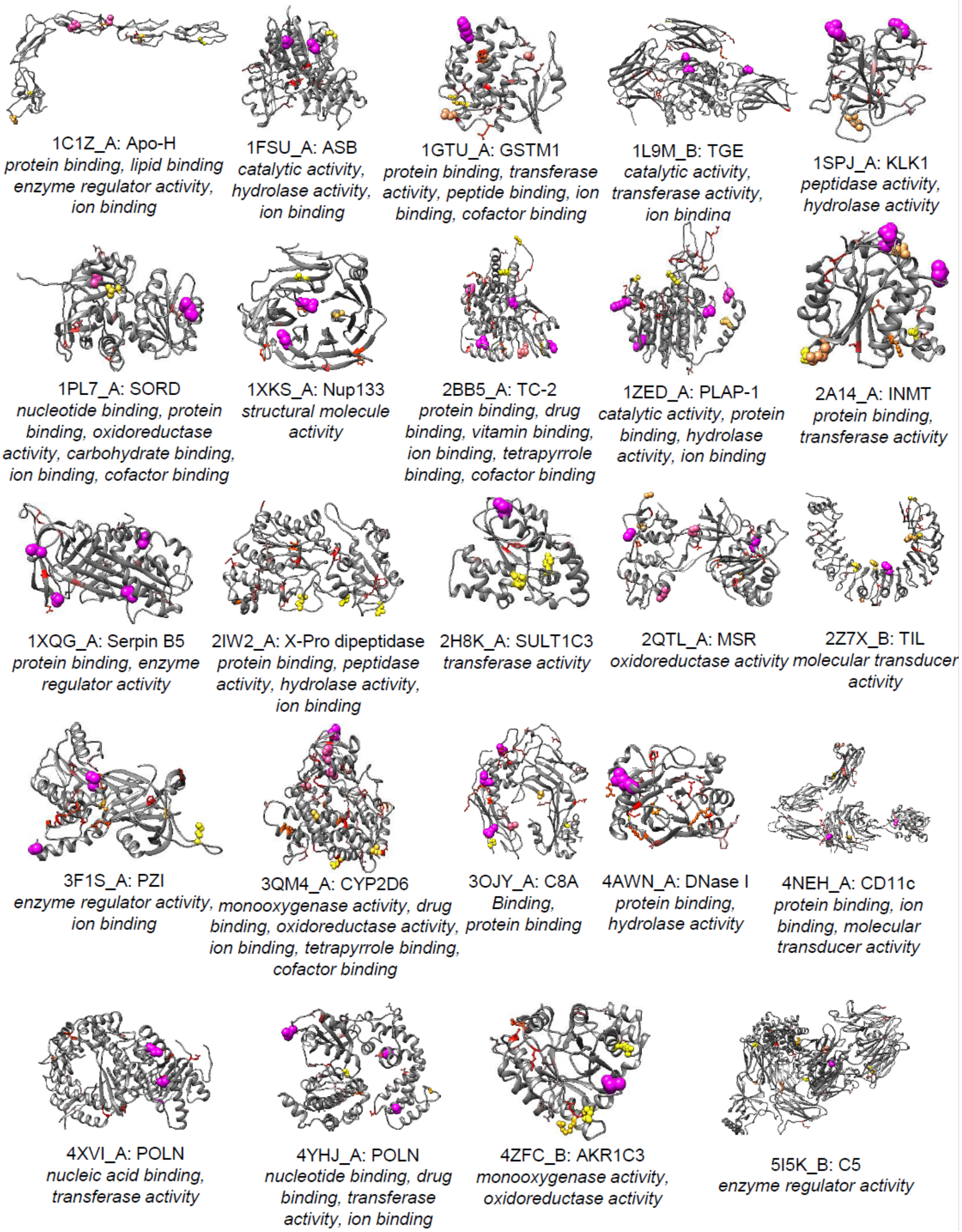


**Figure S4. Highly diverse proteins cover a range of folds and functions.**
A variety of protein structures with high genetic diversity are illustrated along with their gene ontology annotations (molecular function) and 1000G variants. Variant residue color and size indicate frequency across all 1000G populations. Purple spheres (large): ≥ 10% MAF; yellow spheres (small): 1% ≤ MAF < 10%; red sticks: 0.1% ≤ MAF < 1%. Variants with MAF < 0.1% are not included.

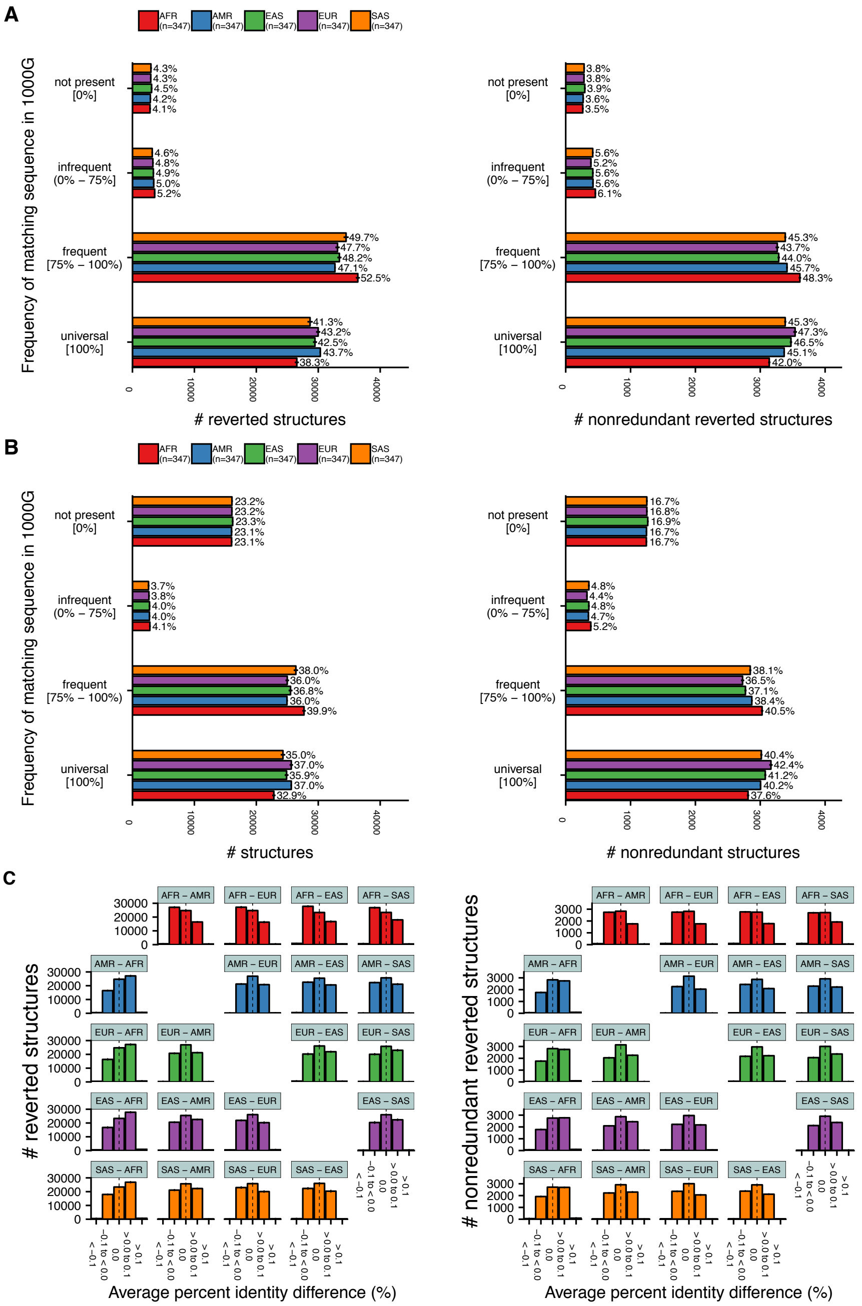


**Figure S5. Sequences from individuals of African ancestry are less represented by and less similar to available protein structures than sequences from individuals from other populations.**

(A) Histograms of the frequency of protein sequences represented by available structures in each population after reverting engineered mutations. Non-AMR populations were down-sampled to match the smaller AMR population (347), and the average over five random samples is presented with standard errors. Histograms based on all chains (left) and only chains within the non-redundant set (right) are included. Note that the difference in fraction matching for AFR and EUR between this figure and Figure 4A is due to different amounts of down-sampling. (B) The same histograms as in (A) but using unaltered raw chain sequences. (C) Differences in average percent identity across all chains (left) and only chains in the non-redundant set (right) for every pair of populations.

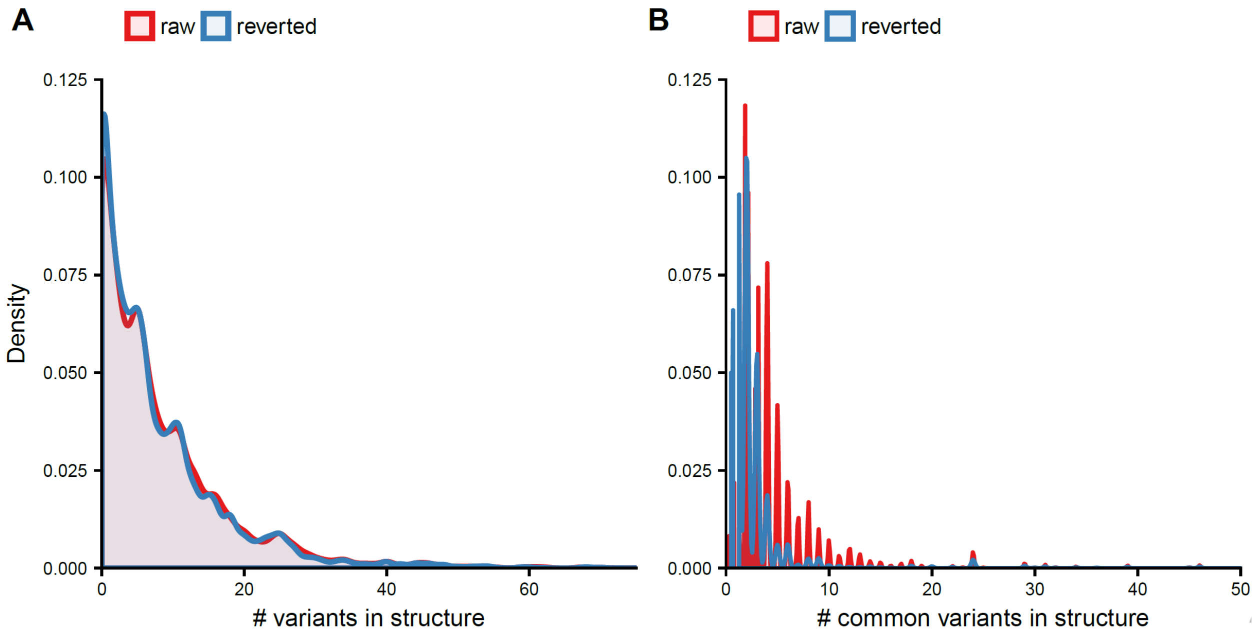


**Figure S6. Sequences differences to available structures are most commonly a few single residue variants.**

(A) Density of the total number of variable amino acids in a protein structure across all structures. Raw structure sequences and those after reverting engineered mutations are overlaid. (B) Density of total number of common variants (frequency in 1000G >= 1%) within a protein structure over all protein structures.

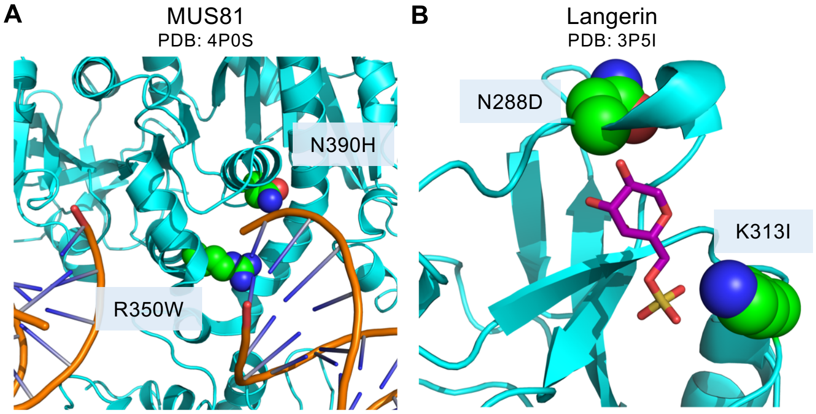


| Variant | AFR | AMR | EAS | EUR | SAS |
|---|---|---|---|---|---|
| R350W | 0.020 | 0.016 | 0.000 | 0.024 | 0.001 |
| N390H | 0.058 | 0.004 | 0.000 | 0.000 | 0.000 |

| Variant | AFR | AMR | EAS | EUR | SAS |
|---|---|---|---|---|---|
| N288D | 0.218 | 0.053 | 0.137 | 0.062 | 0.127 |
| K313I | 0.218 | 0.053 | 0.137 | 0.062 | 0.127 |
| Both | 0.218 | 0.053 | 0.137 | 0.062 | 0.127 |

**Figure S7: Examples of unmodeled variants that influence annotated functional sites.**
(A) Two variants (R350W and N390H) in MUS81, a crossover junction endonuclease, are not represented in the available structures. One variant in particular (N390H) is common in Africans (MAF of 6%) and rarely observed in any other population (EUR MAF of 0%). N390H and R350W both directly modify the 5' end DNA binding pocket that MUS81 forms in complex with EME1 (PDB: 4P0S). (B) Two co-occurring variants (N288D and K313I) disrupt the binding site of terminal 6-sulfated galactose containing oligosaccharides in the protein langerin (CD207, PDB: 3P5I). These variants are at 22% frequency in African populations, but only 6% in European populations. For both proteins, variant frequencies in 1000G populations are listed, and variants are illustrated as atom spheres.

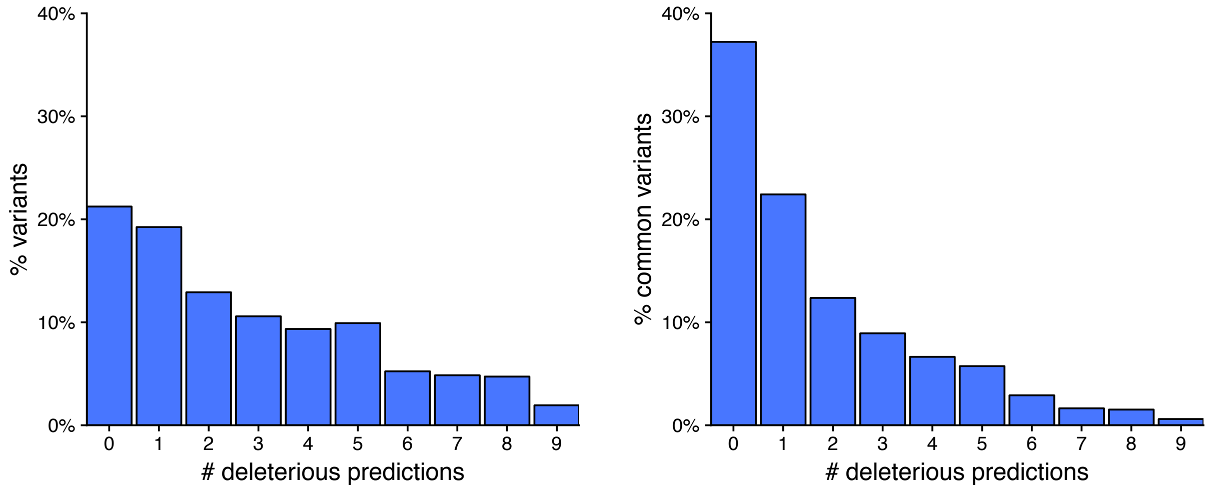


**Figure S8: Percent of variants in 1000G not modeled in structures that are predicted to be functional by nine variant effect predictors.**
The left plot considers all variants unrepresented in structures, and the right plot considers only variants with frequency of at least 1% in any population. Prediction methods include SIFT, PolyPhen2_HDIV, PolyPhen2_HVAR, FATHMM, Mutation Assessor, Meta-SVM, Meta-LR, PROVEAN, and LRT (Methods).

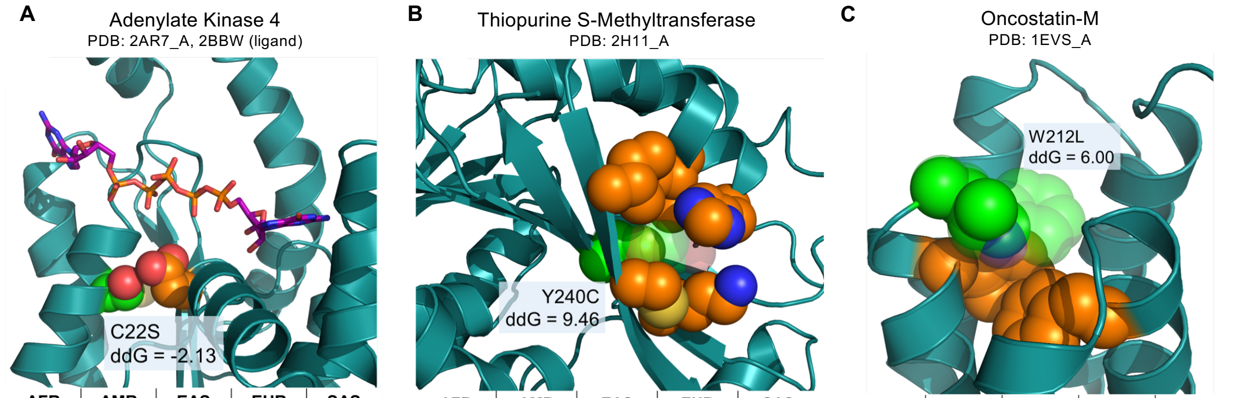


| AFR | AMR | EAS | EUR | SAS |
|---|---|---|---|---|
| 0.508 | 0.072 | 0.273 | 0.018 | 0.074 |

| AFR | AMR | EAS | EUR | SAS |
|---|---|---|---|---|
| 0.067 | 0.058 | 0.022 | 0.029 | 0.017 |

| AFR | AMR | EAS | EUR | SAS |
|---|---|---|---|---|
| 0.000 | 0.000 | 0.014 | 0.000 | 0.0020 |

**Figure S9: Examples of unmodeled variants with different frequencies across populations that influence protein stability.**

(A) Adenylate kinase 4 (AK4) is an enzyme involved in anti-cancer drug sensitivity. The unrepresented S allele at this site is the most common allele in African populations (51%), while it occurs in less than 2% of the European population. This variant is predicted to be significantly stabilizing in PDB structure 2AR7_A. The binding site for diguanosine pentaphosphate (from the superimposed PDB structure 2BBW) is represented by sticks. The atoms of the affected residue are represented as green spheres with the reference amino acid transparent; neighboring residues that interact with the variant residue are shown as orange atom spheres. (B) The common variant Y240C (~4% MAF overall) is predicted to destabilize thiopurine S-methyltransferase (TPMT, PDB: 2H11_A), an enzyme involved in the inactivation of thiopurine drugs in which genetic variation is known to have effects on activity. This variant co-occurs with A154T in patients with TPMT deficiency and has decreased enzyme activity. (C) The variant W212L in oncostatin-M (OSM), a cytokine protein, is common in East Asian populations, but never observed in Africans or Europeans. This variant is predicted to significantly destabilize OSM (PDB: 1EVS_A). This is likely due to W212's role in one of two aromatic stacking groups that make up the protein core..

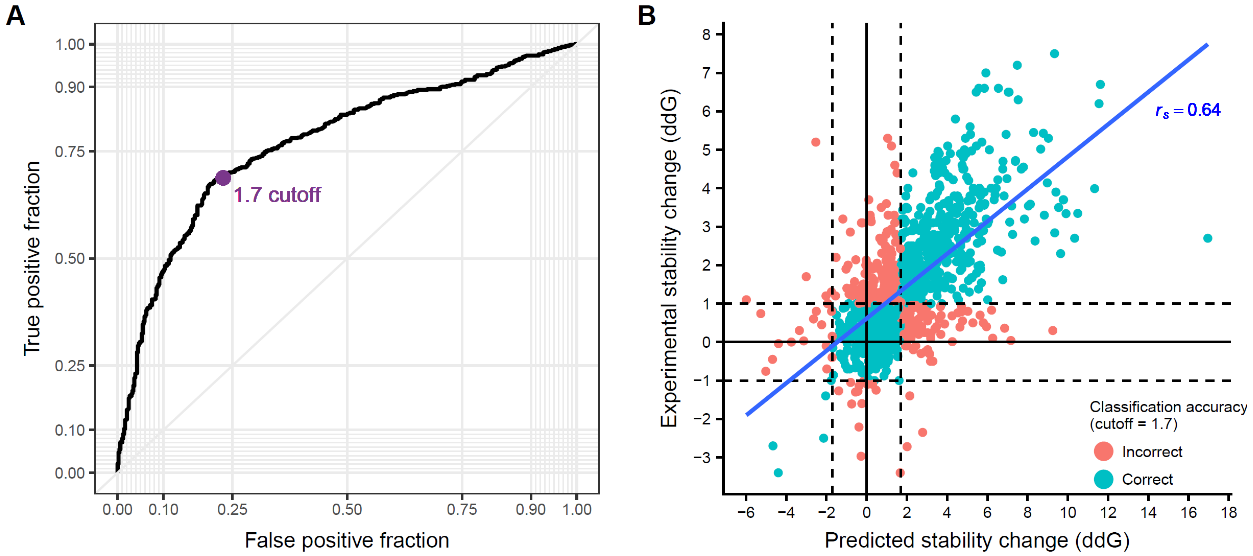


**Figure S10: Benchmarking of Rosetta's ddG monomer stability change predictions.**
(A) Receiver operating characteristic curve of benchmarked fold stability changes with ddG monomer. We selected a cutoff of 1.7 Rosetta Energy Units (purple dot) to optimize the tradeoff between specificity and sensitivity. (B) Correlation of predicted and experimentally determined fold stability changes for the ddG monomer benchmark (Methods). Fit line with Spearman correlation of 0.64 is shown. Each point is a single variant in the benchmark set. Variants are colored according to whether they are correctly categorized as 'stabilizing', 'destabilizing', or 'benign' with the absolute cutoff of 1.7 REU.
